# Refining the transcriptome of the human malaria parasite *Plasmodium falciparum* using amplification-free RNA-seq

**DOI:** 10.1101/852038

**Authors:** Lia Chappell, Philipp Ross, Lindsey Orchard, Thomas D. Otto, Matthew Berriman, Julian C. Rayner, Manuel Llinás

**Affiliations:** Wellcome Sanger Institute, Wellcome Genome Campus, Cambridge CB10 1SA, United Kingdom; Department of Biochemistry & Molecular Biology and Huck Center for Malaria Research, Pennsylvania State University, University Park, PA USA 16802; Department of Biochemistry and Molecular Biology, The University of Chicago, Chicago, IL USA 60637; Institute of Infection, Immunity and Inflammation, MVLS, University of Glasgow, Glasgow G12 8TA United Kingdom; Cambridge Institute for Medical Research, University of Cambridge, Cambridge CB2 0XY, United Kingdom; Department of Chemistry, Pennsylvania State University, University Park, PA USA 16802

## Abstract

*Plasmodium* parasites undergo several major developmental transitions during their complex lifecycle, which are enabled by precisely ordered gene expression programs. Transcriptomes from the 48-hour blood stages of the major human malaria parasite *Plasmodium falciparum* have been described using cDNA microarrays and RNA-seq, but these assays have not always performed well within non-coding regions, where the AT-content is often 90-95%. We developed a directional, amplification-free RNA-seq protocol (DAFT-seq) to reduce bias against AT-rich cDNA, which we have applied to three strains of *P. falciparum* (3D7, HB3 and IT). While strain-specific differences were detected, overall there is strong conservation between the transcriptional profiles. For the 3D7 reference strain, transcription was detected from 89% of the genome, with over 75% of the genome transcribed into mRNAs. These datasets allowed us to refine the 5’ and 3’ untranslated regions (UTRs), which can be variable, long (>1,000 nt), and often overlap those of adjacent transcripts. We also find that transcription from bidirectional promoters frequently results in non-coding, antisense transcripts. By capturing the 5’ ends of mRNAs, we reveal both constant and dynamic use of transcriptional start sites across the intraerythrocytic developmental cycle resulting in an updated view of the *P. falciparum* transcriptome.

## Introduction

There are six species of *Plasmodium* that are known to cause malaria in humans, but most of the estimated 720,000 annual deaths are caused by *Plasmodium falciparum* (1). Although *Plasmodium* spp. have a complex life cycle that involves both invertebrate and vertebrate hosts, it is the asexual development of the parasite in the blood that is responsible for all clinical symptoms of malaria. Blood stage development begins when a newly released, extracellular parasite (a merozoite) invades an erythrocyte, establishing the ring stage of infection, which progresses to the trophozoite stage, during which the infected erythrocyte is extensively modified to enable parasite proliferation (2). The parasite then divides to form a connected group of daughter cells, termed the schizont, which eventually lyses the host erythrocyte, releasing the newly formed merozoites to invade new erythrocytes. Collectively, these steps are known as the intraerythrocytic developmental cycle (IDC), and take 48 hours to complete in *P. falciparum*.

The *P. falciparum* genome is 23.3 Mb in size and encodes over 5,400 genes (3). Most parasite genes are transcriptionally regulated during the IDC, often expressed across multiple time points but with a single peak of maximum abundance per gene (4–6). Another study has compared the IDC transcriptome profiles of three laboratory strains (3D7, HB3 and Dd2; with origins in West Africa, Latin America and Asia, respectively), demonstrating that gene expression was remarkably conserved between strains from across the globe, despite the strains being isolated at different times from disparate geographical locations (7).

These initial analyses of the *P. falciparum* IDC were based on cDNA microarray technology. The first application of RNA-seq to the *P. falciparum* IDC led to alterations in the gene models for over 10% of the ∼5,400 genes, including the identification of 121 new coding sequences (8). This study also confirmed 75% of predicted splice sites and conservatively detected 84 cases of alternative splicing. However, the limitations of available RNA-seq technology at that time meant that the extremely AT-rich UTRs could not be detected on a genome-wide scale; this was probably caused by a combination of difficulties generating AT-rich cDNA and PCR bias against the AT-rich sequences.

The extreme AT-content of the *P. falciparum* genome remains challenging even for current sequencing and alignment technologies; within coding sequences the AT-content is ∼75%, but in non-coding regions the AT-content rises to ∼90-95%. Successive RNA-seq studies (9–16) each used protocols that were not fully optimised for generation of AT-rich cDNA, preventing the full extent of transcription outside of the protein-coding regions of the genome from being captured. One source of bias is random priming of extremely AT-rich RNA fragments; primer binding to these AT-rich fragments is less stable, resulting in fewer AT-rich fragments being converted to cDNA during reverse transcription (17). The use of PCR amplification is another source of bias in RNA-seq, the effect of which is particularly pronounced with AT-rich sequences (18). In whole genome sequencing, the AT-bias can be dramatically reduced with “PCR-free” Illumina sequencing adaptors and omission of PCR amplification steps (Kozarewa et al. 2009). Several of the previous *P. falciparum* RNA-seq studies described thousands of non-coding RNA molecules (ncRNAs) originating from the most AT-rich regions of the genome (López-Barragán et al. 2011; Vignali et al. 2011; Sorber et al. 2011; Hoeijmakers et al. 2013; Siegel et al. 2014; Broadbent et al. 2015; Toenhake et al. 2018), but gaps in sequence coverage in these regions limited assembly of complete transcripts. Many of these predicted ncRNAs have subsequently been discarded during reannotation (19).

To explore the *P. falciparum* transcriptome with minimal bias from the extreme AT-content, we have developed a directional, amplification-free RNA-seq protocol (DAFT-seq) that produces more accurate measures of gene expression. Analysis of the resulting DAFT-seq data revealed extensive transcription between coding regions, particularly of long and often overlapping UTRs. When we applied DAFT-seq to the IDCs of three strains of *P. falciparum*: 3D7 (the genome reference clone, and presumed to be of West African origin) (20), HB3 (a drug sensitive isolate from Honduras) (21) and IT (isolated in Brazil and widely used for studies of antigenic variation) (22), we identified relatively few differences in transcript levels and transcription start sites (TSSs) between these strains. To specifically capture the 5’ ends of mRNAs from the 3D7 strain, we developed a modified amplification-free RNA-seq protocol (5UTR-seq), and confirmed multiple features by sequencing long cDNA molecules from the same clone using the Pacific Biosciences (PacBio) platform. The PacBio platform has been used to sequence *Plasmodium* genomic DNA (23–28), but not yet cDNA. Collectively, these new approaches provide a new view of the *P. falciparum* transcriptome at a greater level of resolution. We provide precise definitions of the boundaries of coding transcripts and comprehensively define 5’ and 3’ UTRs, and TSS positions, on a genome-wide scale. In particular, transcription from bidirectional promoters and overlapping transcripts are common features. These data will be informative for both experimental genetic studies and for further dissecting the mechanisms of transcriptional regulation in *Plasmodium spp*.

## Materials and Methods

### Parasite culture and RNA extraction

*P. falciparum* clone 3D7 was cultured in O+ human erythrocytes and 10% human serum in RPMI-based media, using standard methods (29). RNA extractions used the TRIzol reagent as previously described (30). RNA was quality controlled and quantified using an Agilent Bioanalyzer 2100 Nano RNA chip.

### Directional, Amplification-Free RNA-seq (DAFT-seq) libraries

PolyA+ RNA (mRNA) was selected using magnetic oligo-d(T) beads. Reverse transcription using Superscript II (Life) was primed using oligo d(T) primers, then second strand cDNA synthesis included dUTP. The resulting cDNA was fragmented using a Covaris AFA sonicator. A “with-bead” protocol was used for dA-tailing, end repair and adapter ligation (NEB) using “PCR-free” barcoded sequencing adaptors (Bioo Scientific, similar to Korarewa et al.). After 2 rounds of SPRI cleanup the libraries were eluted in EB buffer and USER enzyme mix (NEB) was used to digest the second strand cDNA, generating directional libraries. The libraries were quantified by qPCR and sequenced on an Illumina HiSeq2000, generating 100bp paired-end reads. Reads were mapped using TopHat2 (31), using directional parameters and a maximum intron size of 5,000 nt.

### 5UTR-seq libraries

PolyA+ RNA was isolated using oligo d(T)-coated magnetic beads. Superscript II reverse transcriptase was used to synthesise first strand cDNA using oligo-d(T) primers and in the presence of template switching oligos, which had the same sequence as those in the Smart-seq2 protocol (32, 33). Template-switching oligos (TSOs) were used to “tag” the end of the cDNA sequences; this tag is used to prime second strand cDNA synthesis. The resulting cDNA was fragmented, made into RNA-seq libraries using an amplification-free protocol based on the DAFT-seq protocol, then were sequenced using 100bp paired-end reads on an Illumina HiSeq2000. 5UTR-seq reads marking the TSS were identified by mapping using SMALT (34) and identified soft-clipped reads containing the TSO sequence. We removed G residues at the end of the TSO sequence that corresponded to the bases in the oligo that were involved in template-switching. We analysed reads in defined windows (up to 2,500 nt) upstream of the annotated translation start site, and limited to finding the single most frequently used TSS at each time point. The threshold using for determining expression was at least 5 reads mapping to the same genomic position. Secondary TSSs were defined to be those with at least half the reads of the main TSS.

### Transcription start site (TSS) clustering and analysis

5UTR-seq alignments were collapsed onto their 5’ ends to define genome-wide CTSSs at each sampled time point. CTSSs were then analyzed using the CAGEr bioconductor package (version 1.12). CTSSs were first normalized and tags-per-million (TPM) were calculated at each position. Positions across the genome where the TPM was greater than 1 were clustered based on distance (20 base-pairs apart or less) into TSS clusters (TCs). Following TSS clustering per time point, each TC was clustered based on distance (100 base-pairs apart or less) into consensus promoter clusters (PCs). These were then filtered by those with an expression level of at least 5 TPM at one time point during the IDC. The resulting PCs were then clustered by their expression profiles using a self-organizing map and predefining the dimensions to 1 by 7. TCs and PCs were annotated using the bedtools closest command. Promoter shape was defined by the interquartile width of each TC as in (35).

### Regulatory element analysis

Protein-binding microarray derived sequence-specific motifs for 24 ApiAP2 domains were obtained from (36) and trimmed to their 6 most informative, consecutive bases or core motifs (see supplementary text). The core motifs were used to search promoters extracted from version 3 of the *Plasmodium falciparum* reference genome using FIMO of the MEME suite and a threshold of 1×10^-2^ (37). Promoters were defined as 1000 bp regions upstream of most frequently used, predicted TSSs.

### Custom exploration of the extended transcriptome

RPKM values were calculated using in-house Perl scripts (Lia Chappell) that use the BEDTools suite (38). Detected of new RNA sequences including UTRs and ncRNAs were identified using a custom approach described further in the see supplementary materials, which also used the BEDTools suite to manipulate blocks of continuous coverage and consider overlaps with other transcriptomic features.

### Detection of splice sites (including) exitrons

A custom script (Lia Chappell) was used to identify spliced reads based on the CIGAR string in the DAFT-seq BAM files. Comparison with existing annotation allowed the detected of previously annotated splice sites. Splice sites in features such as UTRs and ncRNAs was detected by searching for spliced reads overlapping these features. Exitrons were identified by searching for spliced reads mapping within protein-coding exons. At least 5 reads were needed to support detection of each splice site.

### Comparative strain analyses

Detection of expression was defined for a threshold of 5 TPM (tags per million mapped reads, a normalised measure of gene expression). Differential transcript abundance was calculated by first comparing mean Log2(TPM + 1) values across the IDC between strains using the Student’s t-test for each gene. Multiple testing correction was performed using a false discovery rate, as implemented in the qvalue Bioconductor package (version 2.2.2) (39). Differentially abundant genes were identified as having an absolute Log2 fold-change of at least 2 and a q-value less than or equal to 0.05. Correlations between genes were calculated using the Pearson correlation coefficient. Phases were calculated by implementing a non-parametric multidimensional scaling approach as demonstrated in (14) using the MASS R package available through CRAN (version 7.3-51.1).

### Use of published genomics tools

The Artemis genome browser (40, 41) was used to visualise RNA-seq data. Enriched GO terms were identified using the TopGO tool (42). A q-value cutoff of 0.05 was applied.

### Code for processing and analysis of sequence data

Code is available through http://github.com/LiaChappell/DAFT-seq and https://llinaslab.github.io/sanger_rnaseq

## Results

### Directional, amplification-free RNA-seq (DAFT-seq) reveals extensive transcription from the 3D7 genome

We developed an optimised directional, amplification-free RNA-seq protocol (DAFT-seq; **Figure S1**, supplementary materials) that uses adaptors that eliminate the need for PCR (Kozarewa et al. 2009), even for low input quantities of total RNA (≥500 ng). A further critical modification was to synthesise full-length cDNA molecules, which were then fragmented to make libraries; this gave more even coverage in AT-rich regions (**Figure S2**). To map transcripts throughout the IDC, DAFT-seq libraries were generated from seven RNA samples taken from tightly synchronized *P. falciparum* 3D7 parasites at 8-hour intervals from 0 - 48 hours. For most DAFT-seq libraries we obtained around 10 million reads that mapped to the parasite genome (**Table S1**). Mapping of these libraries showed that the majority of each chromosome sequence is transcribed, as shown for chromosome 1 in **Figure 1A**. A striking feature of the data is the extent to which the transcripts extend beyond existing annotated protein-coding exons, defining much larger 5’ and 3’ untranslated regions (UTRs) than previously realised (**Figure 1B,C**).

**Figure 1:**
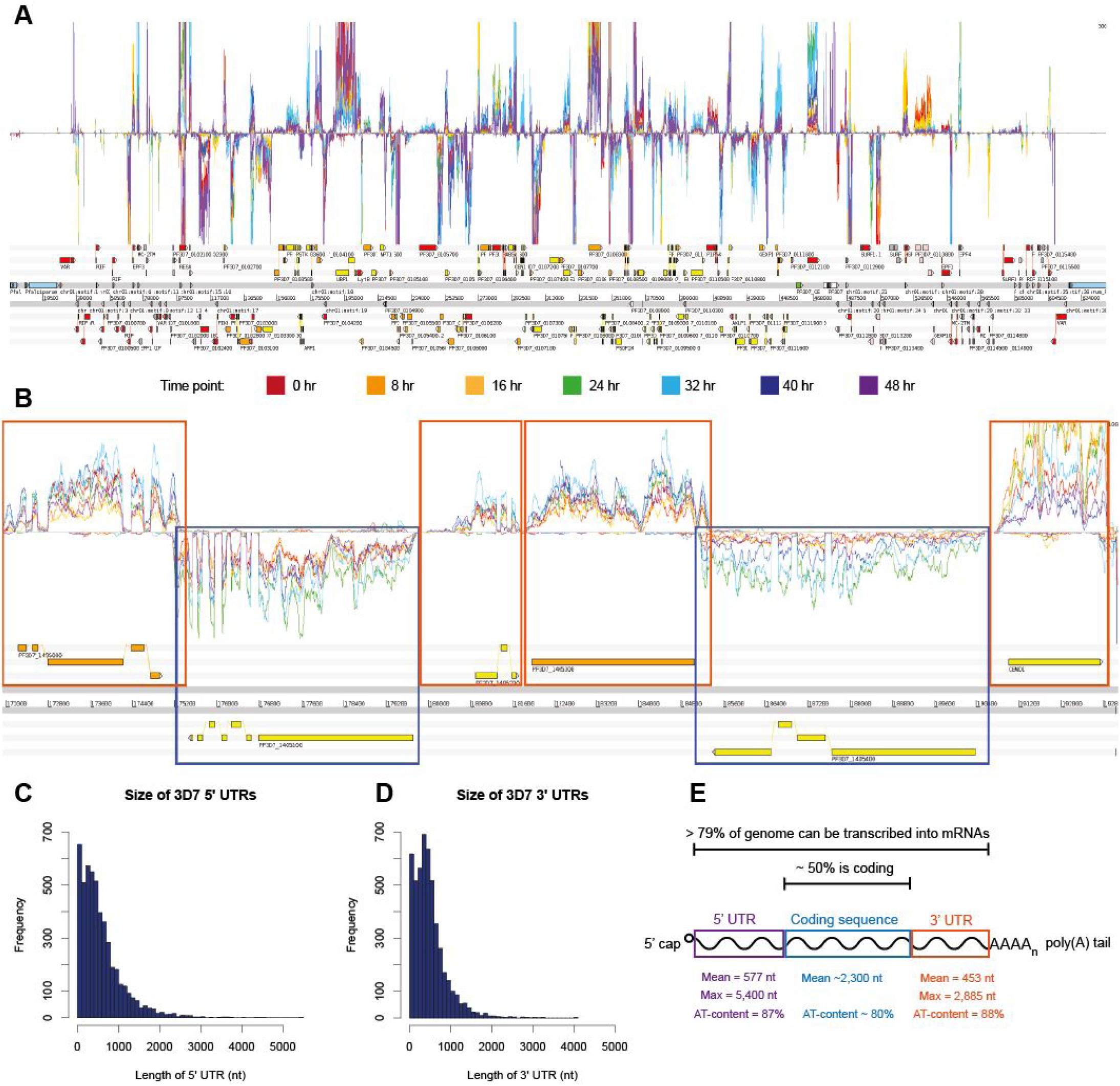
Most of the genome is transcribed as part of mRNAs. A) Overview of DAFT-seq data for the 3D7 time course for all of chromosome 1. Top panel: DAFT-seq coverage for plus strand. Coloured traces represent normalised coverage for each of the seven time points analysed. Middle panel: DAFT-seq coverage for minus strand. Lower panel: annotated gene models for Pf3D7v3 from GeneDB. Legend: colours of coverage traces from each of the seven time points. B) Continuous coverage of DAFT-seq data allows transcript boundaries to be redefined. Orange boxes define boundaries of transcripts on the plus strand and blue boxes define boundaries of transcripts on the minus strand. Colours of coverage traces from the seven time points are the same as those shown above. C) Size of 3D7 5’ UTRs based on continuous coverage of DAFT-seq data. See supplementary information for details of the computational method. D) Size of 3D7 3’ UTRs based on continuous coverage of DAFT-seq data. See supplementary information for details of the computational method. E) Summary statistics to describe the extent of the genome that is transcribed. At least 79% of the genome can be transcribed into mRNA.

We used the DAFT-seq data to define the positions of 5’ UTRs (**Figure S3, Table S2**) for 4,982 genes in the 3D7 genome (94% of those detected as expressed at >5 RPKM in the IDC, **Table S3**, method shown schematically in **Figure S3**). The precise boundary of each UTR likely varies slightly from transcript to transcript, but in order to annotate a single fixed position for each boundary, we estimated the true position by defining the position at which continuous RNA-seq coverage drops below a threshold of 5 reads. We used a more stringent threshold to avoid merging adjacent UTRs on the same strand; the threshold used to define a block of continuous transcription was iteratively increased, in increments of 5 reads. This approach relies on continuous coverage along the length of a transcript, which is a feature and strength of the DAFT-seq protocol. These coverage-based UTRs generally represent the longest 5’ UTR used for the downstream parent gene, although there are some examples where an extremely AT-rich or unmappable sequence produces a break in coverage. Despite the average size of the predicted 5’ UTRs being 577 nucleotides (nt) (**Figure 1C**), we identified several that were extremely long. For example, the start position of the longest 5’ UTR mapped to 5.4 kb upstream of the first protein-coding exon for polyA binding protein (*Pf3D7_1107300*). For some genes (6.5% of genes with detected 5’ UTRs), we found splice sites in their 5’ UTRs, with multiple splice sites detected in some instances.

We also predicted 3’ UTRs for 4,356 genes (**Table S4**), again using the point at which continuous coverage of DAFT-seq data drops below a threshold of 5 reads (82% of genes detected at >5 RPKM in the IDC, **Table S3**). The end of most 3’ UTRs (corresponding to polyadenylation sites) is between 500-1000 nt downstream of the end of the final protein-coding exons (**Figure 1D**), with a mean length of 453 nt. The length of the longest measured 3’ UTR was 2,885 nt downstream of the last protein-coding exon of the glycophorin binding protein (GBP) gene (*Pf3D7_1016300*).

We compared our set of longest observed 5’ UTRs to those annotated by Caro *et al*. (**Figure S4**) and those of Adjalley *et al*. (**Figure S5**). The Caro *et al*. 5’ UTR estimates were on average shorter than our predictions, with mean differences ranging from 7 nt (for time point 4 in the Caro *et al*. data set) to as much as 496 nt (for time point 5). In the Adjalley *et al*. study (43), TSSs were generated using thresholds of varying stringency. Using their most conservative threshold, the mean 5’ UTR length was 213 nt longer than our data set; this may be due to differences in thresholds used in analysis of slightly different data types, as their dataset also contains many TSSs per gene. We also compared our predicted coverage-based 3’ UTRs to those from a previous publication that defined 3’ UTRs by locating mapped reads with non-templated runs of adenines (polyA) (13). Overall, the previous calls were slightly longer on average (523 nt), but covered fewer genes (3,443 genes).

Based on our threshold read depth of >5, we found that 88.5% of the 3D7 genome is transcribed during the *in vitro* IDC. This is higher than previously reported (78%) in a study with 30-fold more reads (∼600M reads) (13), emphasising the value of even coverage in DAFT-seq data, and the fact that evidence for transcription is not simply a function of overall sequencing output. In general, the regions with enhanced DAFT-seq sequence coverage relative to previous datasets were those with the highest AT-content (non-coding regions). For example, down-sampling the reads from the Siegel *et al*. dataset to match the smaller number of DAFT-seq reads in the present study reduced the coverage to 63% of the genome, highlighting that a greater proportion of the transcribed genome is accessible with the DAFT-seq protocol.

We identified continuous blocks of transcription (adjacent transcribed bases in the genome, each with >5 mapped reads) in the DAFT-seq data that overlapped protein-coding genes on the same strand. The blocks of continuous transcription that overlap the boundaries of protein-coding genes cover ∼79% of the genome, and 29% of the genome is transcribed as 5’ and 3’ UTRs of protein-coding transcripts, (summarised in **Figure 1E**). We also find >4% of the genome transcribed from both strands, corresponding to mRNAs on opposite, overlapping strands.

### Properties of Transcription Start Sites (TSSs) in the 3D7 strain

Although the continuous nature of DAFT-seq coverage enabled TSSs to be inferred genome-wide from RNA-seq data alone, we employed two additional strategies to better define the *P. falciparum* UTRs. First, we developed a modified RNA-seq protocol to capture the extreme 5’ ends of capped mRNA transcripts (5UTR-seq) and therefore maximise the signal for defining this region. Like DAFT-seq, 5UTR-seq used PCR-free adaptors to confer the same advantages of accessibility in AT-rich regions of the genome (**Figure S6, S7**). Template-switching oligos (TSOs) similar to those described in the Smart-seq2 protocol (Picelli et al. 2013; Picelli et al. 2014) were used to “tag” the 5’ end of mRNA sequences. We also used PacBio long read sequencing to determine the sequence of long, unfragmented cDNA molecules.

Precise TSSs were located for 3,194 genes (**Table S5)** (67% of those expressed at >5 RPKM in the IDC) using data from all time points. The 5’ UTRs determined based by 5UTR-seq had similar mean length (577 nt) to the 5’ UTRs predicted from DAFT-seq coverage (574 nt; **Figure S8A**). Individual TSSs predicted by the two methods were generally consistent, but there were exceptions. Where the UTR predicted from coverage appeared larger than the 5UTR-seq based prediction, we interpreted this to mean that the coverage-based UTR is the longest possible UTR, while the TSS data represented the most frequently used 5’ UTR. In contrast, a TSS identified by 5UTR-seq that is longer than a coverage-based one was hypothesised to indicate that an extremely AT-rich region had caused a short break in sequence coverage upstream of the gene. Indeed, we observed an enrichment of long homopolymer tracts upstream of 5’ UTRs where coverage breaks occurred (**Figure S9**). Thus we attempted to “repair” gaps in the coverage-based UTRs with the 5UTR-seq predicted UTRs, allowing us to generate a combined “longest observed” UTR set (4499 5’ UTRs, **Figure S9B, Table S6**). These observations were also supported by PacBio reads containing TSOs, where a similar distribution of UTR sizes was observed (**Figure S8C, Table S7**). The longest 5’ UTR detected directly by PacBio was ∼2,600 nt (FT2, *Pf3D7_116500*, **Figure S10A**). We can directly detect the TSS linked with each transcript isoform using this data (**Figure S10B,C**).

A detailed comparison of these multiple data sets for the same genes enables a more nuanced understanding of TSSs, including how they can vary both at a given time point or between time points; **Figure 2A** shows an example of multiple TSSs for the gene encoding glyceraldehyde phosphate dehydrogenase (GAPDH, *Pf3D7_1462800*). We found that 90% of TSSs fell outside both annotated exons and introns, 9% fall within annotated exons, and 1% fall within annotated introns (**Figure 2B**). This approach relied on prior knowledge of the position of start codons, and was limited to a maximum window of 2,000 nt upstream of the start codon. Finally, to determine whether TSSs are constant or dynamic, we identified 422 genes **(Table S8)** with sufficient coverage depth (>5 reads per TSS per time point) to independently call TSS-based 5’ UTRs at each of the seven time points from the 3D7 IDC. We found that 55% (232 genes) of these genes showed the same major TSS throughout the IDC, such as the gene encoding the 40S ribosomal protein S3 (*Pf3D7_1465900*). However, a number of genes showed distinct temporal changes of TSS usage throughout the IDC, including GAPDH (*Pf3D7_1462800*; **Figure 2A**), where the distribution of TSS peaks shifts closer to the coding sequence (CDS) at the time point associated with peak mRNA levels. The converse trend is seen for the knob-associated histidine-rich protein gene (KAHRP, *Pf3D7_0202000*, **Figure S11**), revealing dynamic TSS use throughout the IDC.

**Figure 2:**
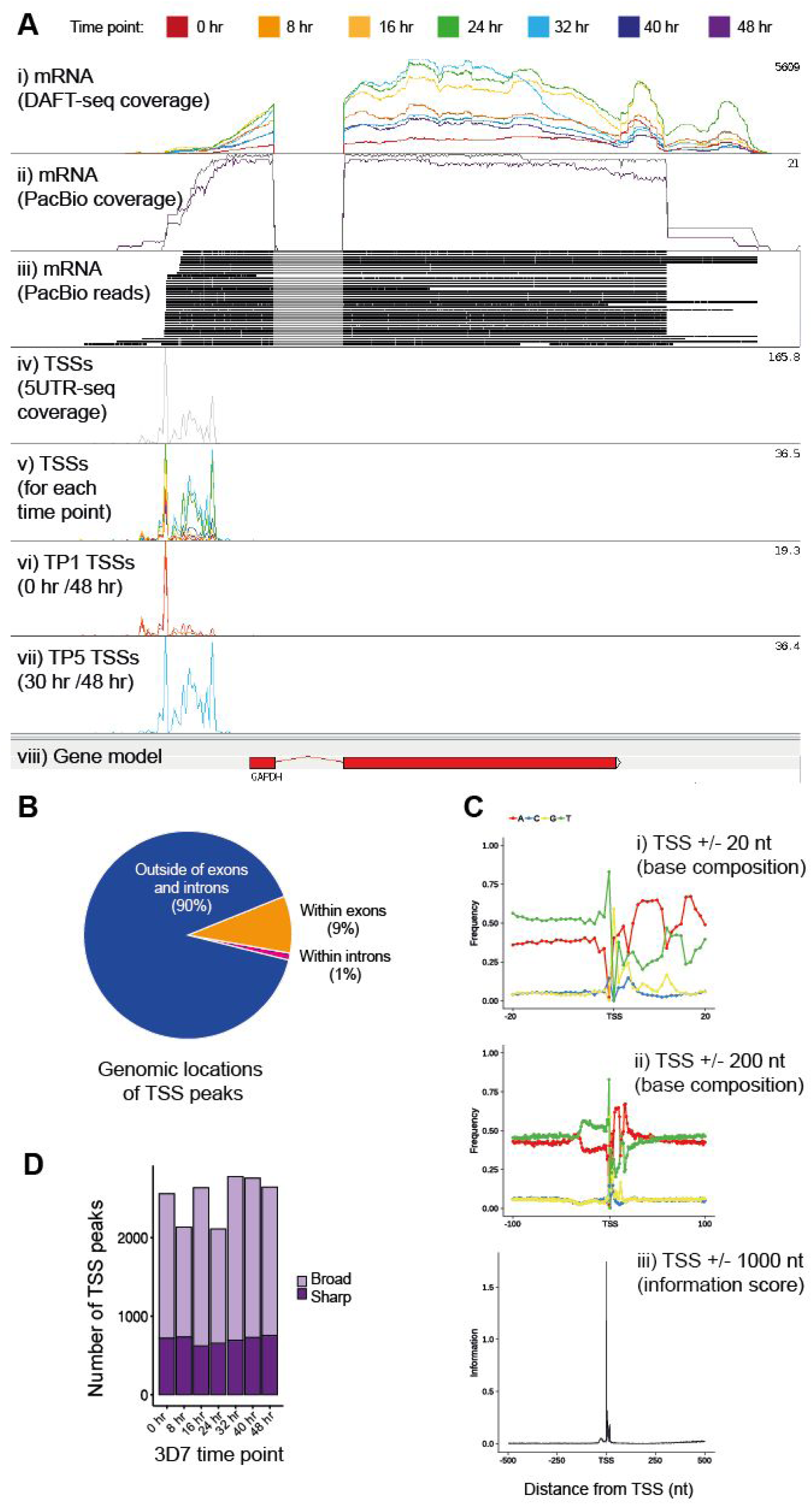
Properties of TSSs and promoters. A) Different library types show different properties of 5’ UTRs and TSSs for the gene encoding GAPDH (*Pf3D7_1462800)*. DAFT-seq coverage (i) can be used to determine the longest possible 5’ UTR. Long read sequencing with PacBio (ii, iii) can be used to directly link a specific TSS with the rest of the transcript structure. Direct detection of TSSs with 5UTR-seq data (iv) reveals a range of different TSSs, which have different prevalences at different time points (v-vii). The first track (i) illustrates 7 DAFT-seq libraries, showing continuous coverage along the length of the gene, and variable steady state levels of mRNA throughout the time course. The next two tracks show PacBio coverage (ii) and reads (iii); these long reads can link variation in the TSS to the structure of the rest of the transcript. The fourth track (iv) shows the extreme 5’ end of mRNAs detected with all of the 5UTR-seq data. This data can be separated by time point (track v), with examination of individual time points showing that the most common TSS early in the time course (track vi) is further upstream from the coding sequence than the most common TSSs later in the time course (track vii). B) Genomic locations of the TSS peaks identified using 5UTR-seq data. The vast majority of the TSS peaks in this data set (90%) fell outside of annotated exons and introns. A small proportion (9%) were within exons, while 1% were within introns. C) Patterns in the base composition around TSSs were identified using the precise TSS positions inferred from the 5UTR-seq data. Windows are shown for a 20 nt distance (i) and a 100 nt distance (ii). Calculation of the information content of the base composition for a 1000 nt window shows that it peak around the inferred TSS. D) Number of TSS peaks in broad or sharp categories for each of the seven time points in the 3D7 time course.

### Sequence features and genomic location of *P. falciparum* TSSs

The information in the 5UTR-seq dataset enabled us to quantify variation in position and time of TSSs associated with the same gene, but our initial analysis was limited by requiring prior knowledge of gene annotation. To address this, the CAGEr package (44) was used to cluster 5UTR-seq reads independently of gene annotation in two stages. First “tag clusters” (TCs) were formed from reads that mapped within 20 nt of each other, for each of the time points (**Table S9**). Depending on the time point, 89-94% of these TCs mapped outside annotated coding and intronic regions. Unlike the comparison between UTR positions for annotated genes, the genome-wide locations of the 5UTR-seq TSSs differ significantly with those in another recent study that tags the 5’ ends of mRNAs using a different approach (43). The authors reported that 49% of all “TSSs blocks” were downstream of the annotated start codons.

Next, nearby TCs (within 100 nt of each other, independent of time point) were grouped into “promoter clusters” (PCs), for each gene. At different time points, we found as many as 37-45% of genes had multiple annotated TCs. While most genes appeared to use a single TSS per time point, our data suggest that some use as many as 14 (**Table S10** and **Figure S12**). We calculated the nucleotide frequency in the regions surrounding the TSS-based 5’ UTRs. We found a global trend for enrichment of thymine residues upstream of TSSs, followed by an enrichment of adenine residues downstream of TSSs; this can be seen both locally (+/- 20 nt) and at greater distances (+/- 1000 nt; **Figure 2C**). In addition, we find that transcription preferentially starts with a pyrimidine-purine dinucleotide, the most preferred being TG. These features are similar to those of highly expressed TSSs described for yeast, mouse, and human (45, 46), suggesting that this is a general feature of promoters that is conserved across a broad range of eukaryotes. We also found that deviations from the average base composition were localised to the site of the TSS itself and not beyond (**Figure 2C**). This signature was also seen for TSSs predicted within introns and exons, albeit with a much weaker signal-to-noise ratio (**Figure S13**).

While most genes contain a primary TSS, we also identified 2,157 genes with a strong distinct secondary TSS in the 5UTR-seq data set (**Table S11,** 68% of genes of the original 3,194). This observation suggests that the potential model of a single “sharp” TSS per gene is inadequate to explain the landscape of transcription initiation in *P. falciparum*. To analyse the proportion of TCs that could be categorised as “sharp” or “broad” we used TCs categorised by their interquartile width (44). In other eukaryotes sharp promoters are often associated with initiation by RNA pol II protein complexes, whereas broad promoters are associated with activation by CpG islands in metazoans, and potentially other mechanisms linked to maintenance of an open chromatin state (45). Analysing each time point separately, we found that many promoters (33-39%, depending on the time point) had a broad shape, establishing that *P. falciparum* transcription initiation is more variable than previously recognised (**Table S9**, **Figure 2D**). Interestingly, the expression level from broad promoters was greater on average than that from sharp promoters, which is inconsistent with results found in other organisms and suggestive of a functional distinction between these two types of promoters (**Figure S13**). Despite these observations, sequence features were highly similar for both types of promoters (**Figure S14**). Additional studies will be required to address whether these differences are of functional relevance.

### Transcription factor (TF) binding motifs relate to TSSs

Using our newly defined 5’ UTRs, we compared the relative location of all known DNA binding motifs associated with the Apicomplexan AP2 (ApiAP2) family of DNA binding proteins, which are considered the major sequence-specific transcriptional regulators present in *Plasmodium* parasites (47–49). Previous studies predicted known ApiAP2 DNA-binding motifs within a defined distance (1-2 kb) upstream of start codons, correlating possible binding sites with the peak time of expression of the ApiAP2 proteins and their putative target genes (36, 50, 51). We remapped all known ApiAP2 binding sequences to the 3D7 genome **(Table S12)** and selected motifs up to 1000 nt upstream of annotated start codons **(Table S13) and** the most frequently used TSSs **(Table S14)**. Using the new TSSs significantly reduced the putative targeted binding motifs genome-wide by 33.8% and sequence search space by 38.3%. While not statistically significant, we found that motif occurrences were biased within 250 bp upstream of predicted TSSs; we speculate that this could be a feature of gene regulation within a compact genome (**Figure S15**). For genes with multiple predicted TSSs at different IDC timepoints, such as *kahrp* (**Figure S16**), independent motif searches were performed for each TSS. While overlapping and distinct motifs were observed for the long and short isoforms, future experiments will be required to functionally validate the differential role of these isoforms.

### Expression of adjacent gene pairs

By improving the accuracy of UTR predictions, the extent to which the *Plasmodium* parasite uses bidirectional promoters became strikingly apparent (**Figure 3A**). Bidirectional promoters have been described in multiple species, including human (52, 53) and yeast (54), where they regulate up to half of all protein-coding transcripts. In metazoans, a distance of less than 1000 bp between head-to-head genes has previously been used to define bidirectional promoters (52, 53). In the *P. falciparum* 3D7 genome there are 1,492 pairs of protein-coding genes in a “head-to-head” orientation (**Table S15**). Few gene pairs have overlapping 5’ UTRs (<0 nt of sequence between TSSs), and when they do overlap, the overlap is small. The median distance between the 5’ UTRs for most pairs of head-to-head genes is 548 nt. In general, we observed positive correlations in the gene expression patterns of head-to-head gene pairs where the distance between the 5’ UTRs was less than 1,000 nt (**Figure 3B**). For example, the start codons for heme oxygenase (*Pf3D7_1011900*) and a putative RING zinc finger protein (*Pf3D7_1012000*) are 1499 bp apart, but their 5’ UTRs are separated by only 298 bp and their expression patterns are highly correlated (**Figure 3C**). Therefore these two genes are likely to share a single bidirectional promoter. We used *in silico* predicted ApiAP2 TF DNA binding motifs, the genetic sequence for this newly defined promoter, and genome-wide gene expression profiles of the genes to predict that the likely ApiAP2 regulators of this promoter are encoded by the genes *Pf3D7_0730300* and *Pf3D7_1305200*. In total, we identified 401 pairs of genes that are potentially regulated by bidirectional promoters in *P. falciparum*, using an expression correlation cutoff of R > 0.5 and a distance between the 5’ UTRs of less than 1,000 nt (**Table S16**).

**Figure 3:**
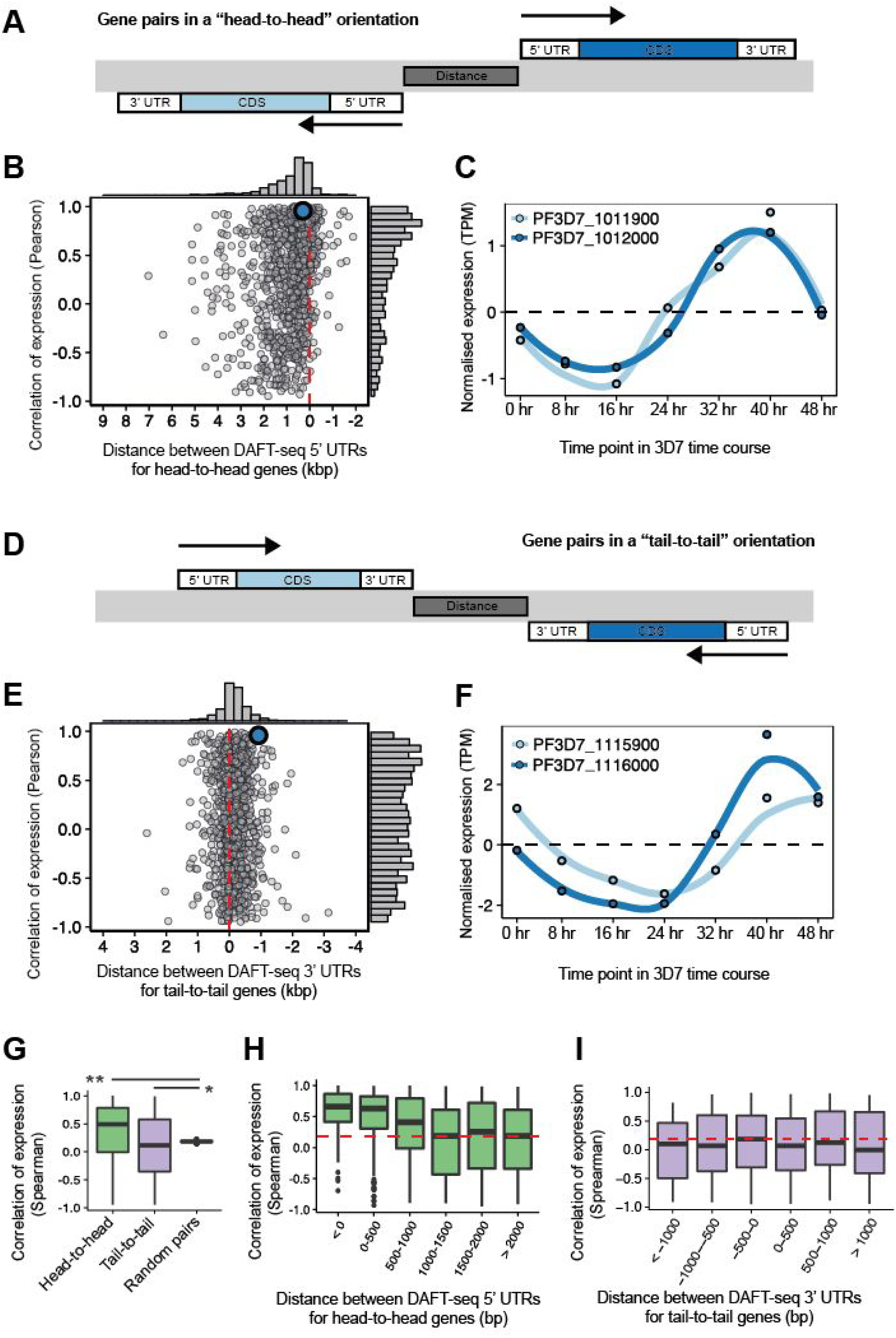
Expression patterns of adjacent transcripts. A) Schematic diagram of gene pairs in a “head-to-head” orientation (also known as divergent gene pairs). The black arrows represent the direction of transcription and the dark grey box between the genes represents the intervening genomic sequence that is between the longest detection version of both 5’ UTRs. B) Correlation of gene expression (in TPM, using Pearson correlation) for 1119 pairs of head-to-head genes with annotated 5’ UTRs plotted against the distance of intervening genomic sequence. The median intervening sequence length is 548 bp (without annotation of 5’ UTRs this distance was 1946 bp). The median correlation of expression was 0.49, with the distribution showing a positive skew. An individual region (blue) is shown in more detail in panel C. C) Expression profiles through the 3D7 time course for a head-to-head gene pair (*Pf3D7_1011900*, heme oxygenase and *Pf3D7_1012000*, RING zinc finger protein, putative). The steady state levels of these two genes is tightly correlated, with an R value of 0.96 (measured by spearman correlation). D) Schematic diagram of gene pairs in a “tail-to-tail” orientation (also known as convergent gene pairs). The black arrows represent the direction of transcription and the dark grey box between the genes represents the genomic sequence that is between the longest detection version of both 3’ UTRs. E) Correlation of gene expression (in TPM, using Pearson) for 1059 pairs of tail-to-tail genes with annotated 5’ UTRs plotted against the distance of intervening genomic sequence. The median intervening sequence length is −124 bp, i.e. most 3’ UTR pairs overlap (without annotation of 3’ UTRs this distance was +657 bp). The distribution of correlation values includes both strongly negative and strongly positive relationships. An individual region (blue) is shown in more detail in panel F. F) Expression profiles through the 3D7 time course for a tail-to-tail gene pair (*Pf3D7_1115900*, DHHC9 and *Pf3D7_1116000*, RON4). Despite an overlap of 327 nt in the 3’ UTRs the steady state level of these genes is strongly correlated, with a spearman correlation of value 0.81. G) Correlation of expression profiles (Spearman) of 1000 neighbouring gene pairs for head-to-head, tail-to-tail and randomly selected gene pairs. The 1000 random neighboring gene pairs were randomly sampled 1000 times from all annotated head-to-head, tail-to-tail and tandem gene pairs. Mean correlations were 0.35, 0.10, and 0.18 for head-to-head, tail-to-tail and random orientations, respectively. Wilcoxon rank sum test was used to determine significance between groups. P-values of 2.2e-16 and 3.8e-3 when comparing head-to-head and tail-to-tail groups to random pairings, respectively. H) Correlation of expression profiles (Spearman) neighbouring gene pairs for head-to-head gene pairs, binned by intervening genomic distance. I) Correlation of expression profiles Spearman) neighbouring gene pairs for tail-to-tail gene pairs, binned by intervening genomic distance.

The *P. falciparum* 3D7 genome also contains 1,503 pairs of genes arranged in a “tail-to-tail” orientation (**Table S17**). We found that the 3’ UTRs overlap for many of these tail-to-tail gene pairs (**Figure 3D**). In total, we identified 1,036 genes (518 pairs of genes) where the 3’ UTRs overlap by at least 100 nt. Longer 3’ UTR overlaps were common; we detected 136 genes (68 pairs of genes) where the overlap was greater than 500 nt and 317 3’ UTRs which overlap the protein-coding sequence of the neighbouring gene. The largest 3’ UTR overlap was 1692 nt and is between genes encoding an acyl-CoA synthetase and a protein kinase (*Pf3D7_1238800* and *Pf3D7_1238900*, respectively). In an extreme example, 2,432 nt of the protein-coding region of *Pf3D7_1244400* overlaps with the 3’ UTR of the adjacent gene (*Pf3D7_1244500).* We expected to find negative correlation for gene pairs with overlapping 3’ ends, due to the possibility of transcriptional interference, in which collision of convergently transcribing RNA polymerases leads to premature termination of transcription (55, 56). However, in contrast to 5’ UTRs (**Figure 3C**), we found little correlation between overlapping 3’ UTR distance and gene expression profiles of tail-to-tail gene pairs (**Figure 3E**). To rigorously determine if the different classes of adjacent gene pairs are coregulated, we further determined the patterns of correlations between 1000 pairs of head-to-head, tail-to-tail and randomly selected gene pairs (**Figure 3G**). The mean correlations were 0.35, 0.10 and 0.18, respectively, consistent with an important role for bidirectional promoters in regulating head-to-head genes in *P. falciparum*. We further binned these correlations by distance for head-to-head (**Figure 3H**) and tail-to-tail genes (**Figure 3I**). Correlation of expression is inversely related to distance for head-to-head genes, which is consistent with the presence of bidirectional promoters present in the most closely adjacent 5’ UTRs. While the average correlation between tail-to-tail gene pairs was also positive, it was significantly less than randomly selected gene pairs suggesting that transcriptional interference may indeed be playing a role, albeit perhaps not a significant one.

### Non-coding transcripts associated with the 5’ end of mRNAs

We also identified a second form of correlated bidirectional promoters in the 3D7 genome between pairs of coding mRNAs and non-coding transcripts (ncRNAs). In several instances we could detect transcription upstream of the 5’ UTRs of mRNA transcripts on the strand opposite to that of the coding mRNA. The expression pattern of these ncRNAs are strongly correlated with the temporal expression of the adjacent coding mRNA, suggesting that these transcript pairs are co-regulated (shown schematically in **Figure 4A)**. These non-coding transcripts were identified for genes where there is several kb of non-coding sequence upstream of the CDS (**Figure 4B**), where there are multiple genes on the same strand (**Figure S17A**) and for genes in a head-to-head orientation (**Figure S17B**). Therefore, the most likely explanation for the observed coregulation is bidirectional transcriptional activity from the TSS that is associated with the mRNA, as has been reported in other eukaryotic species (57, 58). To determine the genome-wide prevalence of this feature, we determined the correlation of transcript levels between all pairs of sense mRNAs and putative antisense ncRNA transcripts using a 2000 nt region upstream of the longest detected 5’ UTR (from our coverage-based 5’ UTRs). In total, we found 337 pairs of “transcriptionally linked” mRNA/ncRNA transcript pairs throughout the 3D7 genome (**Table S18**), which we provisionally describe as “TSS-associated RNAs” (TSSa-RNAs), similar to those previously described in mouse ES cells (58). The functional role of these TSSa-RNAs remains to be determined.

**Figure 4:**
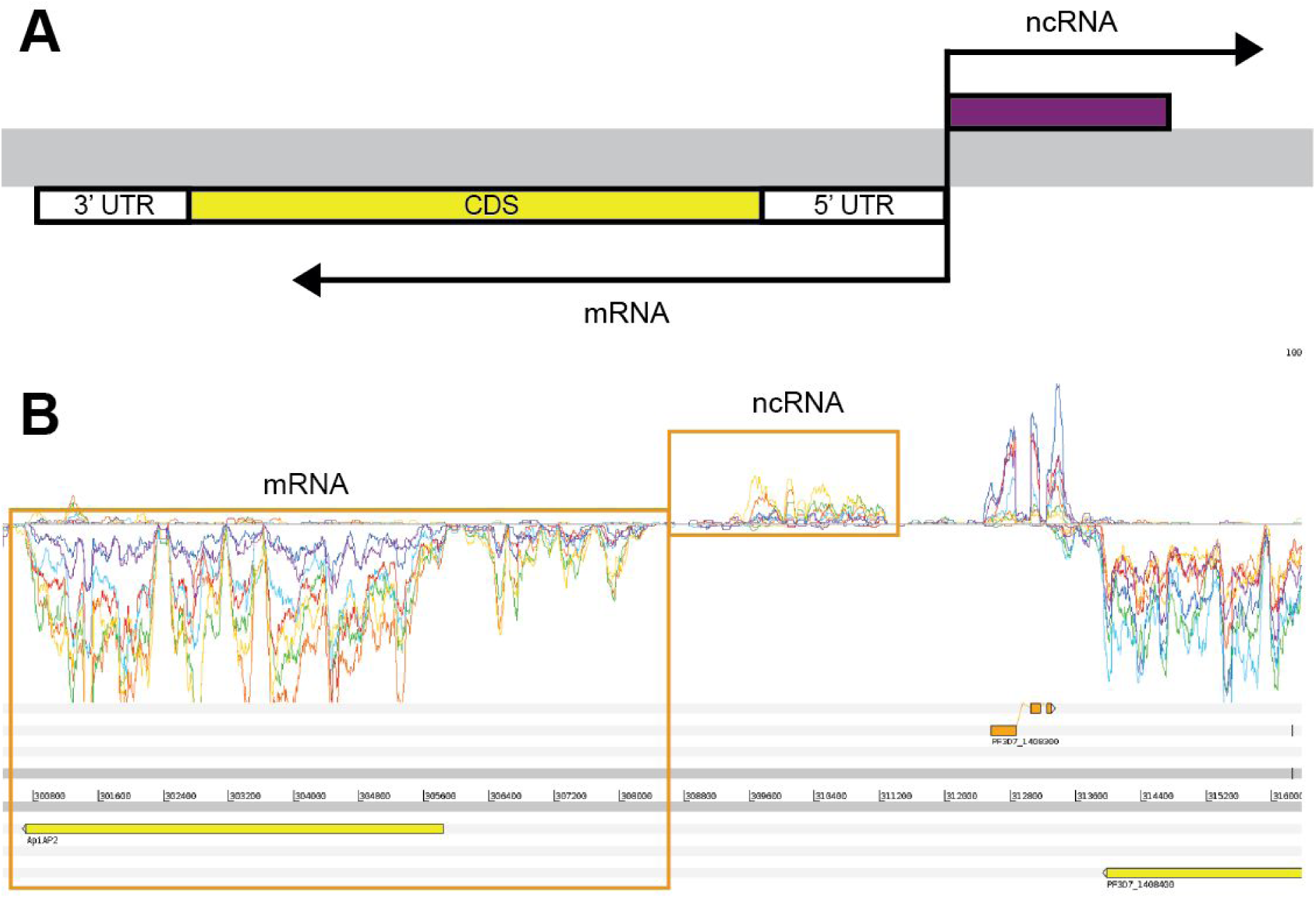
Non-coding RNAs that may share promoters with nearby mRNAs. A) Schematic showing the relative orientations of ncRNAs that may share a promoter sequence with an adjacent mRNA. Black arrows show the direction of transcription. The pair of transcripts may share a bidirectional promoter. B) Example of a ncRNA and an mRNA (*Pf3D7_1408200*, which encodes an ApiAP2 protein) that may share a bidirectional promoter. The transcript pair are shown in the same orientation as the diagram in the top panel. The expression profiles of the two transcripts are correlated through the 3D7 time course.

Other studies (9, 13, 14) have described thousands of ncRNA fragments mapping to the 3D7 genome, although a recent study has called many of these into question (Böehme et al 2019). Because the reference genome has not been annotated with these predictions in online databases, comparisons to these ncRNA is difficult. However, based on our new evidence we suggest that many, if not the majority, of these previously predicted ncRNAs can be considered “orphan fragments” of UTRs, while others represent TSS-aRNAs. However, we also find evidence of independent ncRNAs in our dataset, such as the ncRNA shown in **Figure S18**. After removing fragments of ncRNA which we suspect are part of annotated features, 5’ UTRs, 3’ UTRs, TSS-aRNAs, we find that 1.9% of the genome sequence has transcriptional activity consistent with currently unannotated ncRNA features (**Table S19**).

### Novel splice sites included exitrons (exonic introns)

To identify additional functional features within our extended mRNA transcript models, we examined all spliced reads in the DAFT-seq data sets to annotate splice sites across the 3D7 transcriptome. Splice site predictions were further categorised based on overlap with features such as previously annotated introns or predicted UTRs (**Figure 5A**, **Table S20**). We find that most introns are smaller than 200 nt (**Figure 5B)**. We also evaluated the prevalence of alternative splicing, and found that 6.9% of genes (365 genes) had alternative splice forms within their protein-coding regions **(Table S21)**. Previous studies reported alternative splice forms in 1.5% of detected genes (8) and 4.5% of detected genes (11). Figure **S19** shows two alternative isoforms of the gene *Pf3D7_0316300* are captured in PacBio reads, where the reads capture different TSSs and different use of splice sites.

**Figure 5:**
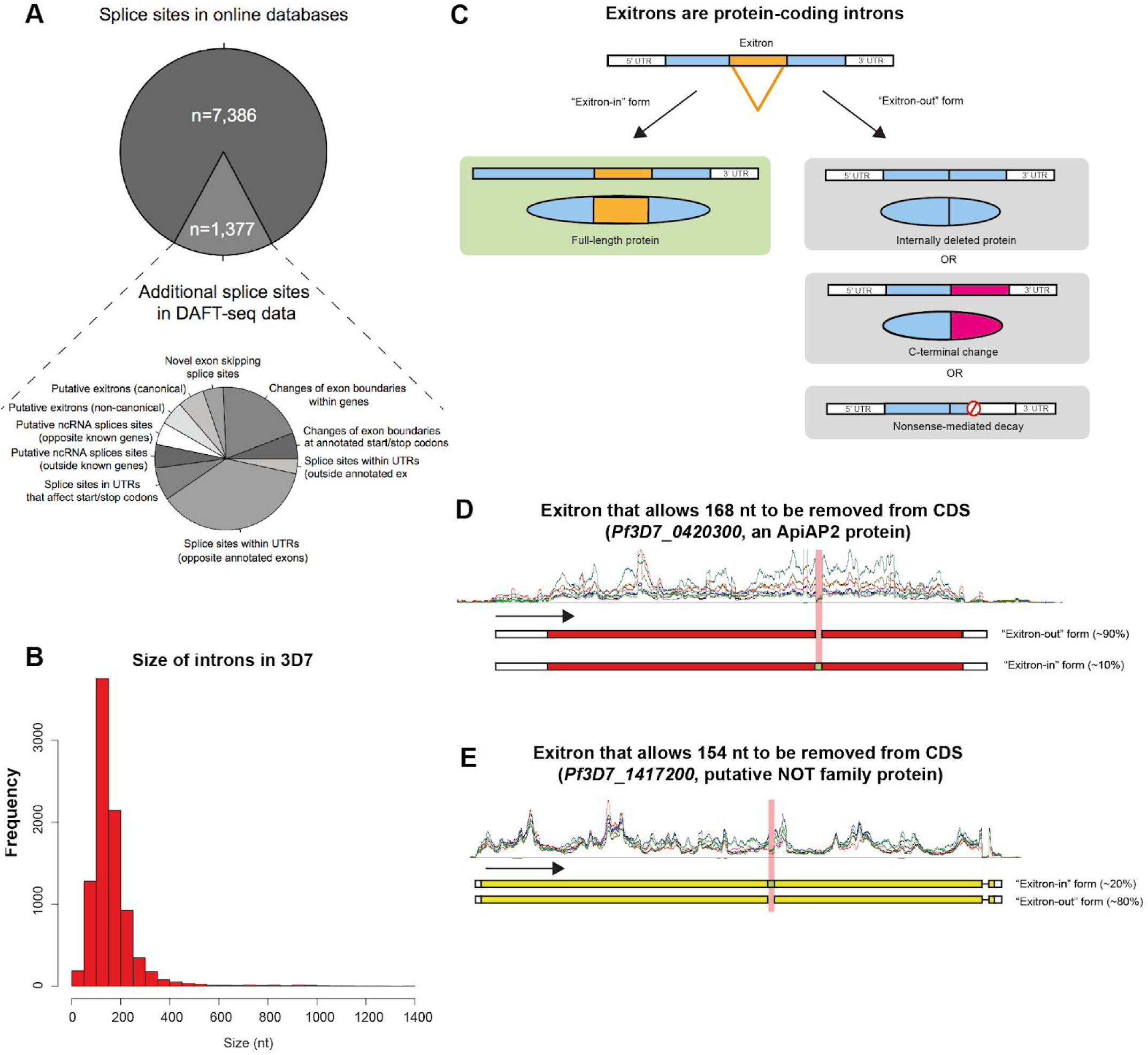
Novel and updated features within genes. A) Splice sites detected in the 3D7 time course using DAFT-seq data. A total of 8,206 splice sites were supported by at least 5 reads; of these, 7,386 were in online databases. There are 1,377 splice sites not described in online databases (17% of the total), including 659 in UTRs. B) Length of introns found in this study, using the DAFT-seq data for the 3D7 time course. C) Schematic diagram of “exitrons” (protein-coding introns) modified from Marquez et al (2015). An exitron may be retained in an mRNA, generating a fully coding mRNA sequence (left). If the excised sequence is a multiple of 3 nt the open reading frame is maintained and a full-length protein may be produced (top right). If the excised sequence is not a multiple of 3 nt this may generate a change to the C-terminus (middle left) or cause the transcript to be degraded through the nonsense-mediated decay pathway (bottom right). D) Exitrons are present in the 3D7 transcriptome. There are examples of mRNAs where a multiple of 3 nt are excised, such as *Pf3D7_0420300* (which encodes an ApiAP2 protein). Here ∼90% of the reads support the “exitron-out” form, with ∼10% of the reads supporting the “exitron-in” form. E) Exitrons are present in the 3D7 transcriptome. There are examples of mRNAs where even numbers of nt are excised, such as *Pf3D7_1417200* (which includes a putative NOT family protein). For this gene ∼80% of the reads support the exitron-out form, with ∼20% of the reads supporting the “exitron-in” form.

Exitrons (exonic introns) are intron-like features that can be found in protein-coding sequences whose retention or loss facilitates an additional level of regulatory control and proteome complexity (**Figure 5C**). To date, exitrons have been reported only in *Arabidopsis thaliana* and human tissues (Marquez et al. 2015). Importantly, splicing of exitrons does not always retain the continuity of the protein open reading frame, resulting in protein diversification. We detected 155 splicing events in annotated protein-coding sequences that are consistent with exitrons (**Table S22**). For example, for *Pf3D7_0420300*, an ApiAP2 gene, 168 nt (a multiple of 3 that retains the open-reading frame) is spliced out in ∼90% of the detected transcripts (**Figure 5D**). However, ∼10% of the transcripts retain the intron, which has an open reading frame that is fully coding. In another gene, *Pf3D7_1417200*, the CDS is interrupted (80%) by an intron that is protein-coding and 154 nt in length (**Figure 5E**). Excision of this exitron disrupts the coding sequence of the downstream protein as the reading frame is not conserved. Other exitrons we found contained non-canonical splice sites and were located in genes encoding exported proteins that may interact with the host, such as glycophorin binding protein (*Pf3D7_1016300*). The predicted exitrons were also detected in the PacBio long read sequencing data (an example is shown in **Figure S20**).

### Comparing DAFT-seq transcriptomes for multiple *P. falciparum* strains

To define whether the extensive transcription found in 3D7 is unique to this strain, or reflects general features of *P. falciparum* biology, we measured the DAFT-seq IDC transcriptomes of the HB3 and IT strains and compared them to 3D7 by mapping the reads to the 3D7 reference genome. DAFT-seq expression data from all three strains was highly correlated to existing high-resolution hourly 3D7 microarray data, as expected (7) (**Figure S21**). We determined whether gene expression could be detected above a threshold of 5 TPM (tags per million mapped reads, a normalised measure of gene expression to best compare between samples) for each gene in HB3, IT and 3D7. Given that many genes within the subtelomeric regions are highly polymorphic, these regions were excluded to avoid mapping issues. This resulted in a common set of 4,371 “core” genes (**Figure 6A**) that were expressed by all strains (**Table S23**).

**Figure 6.**
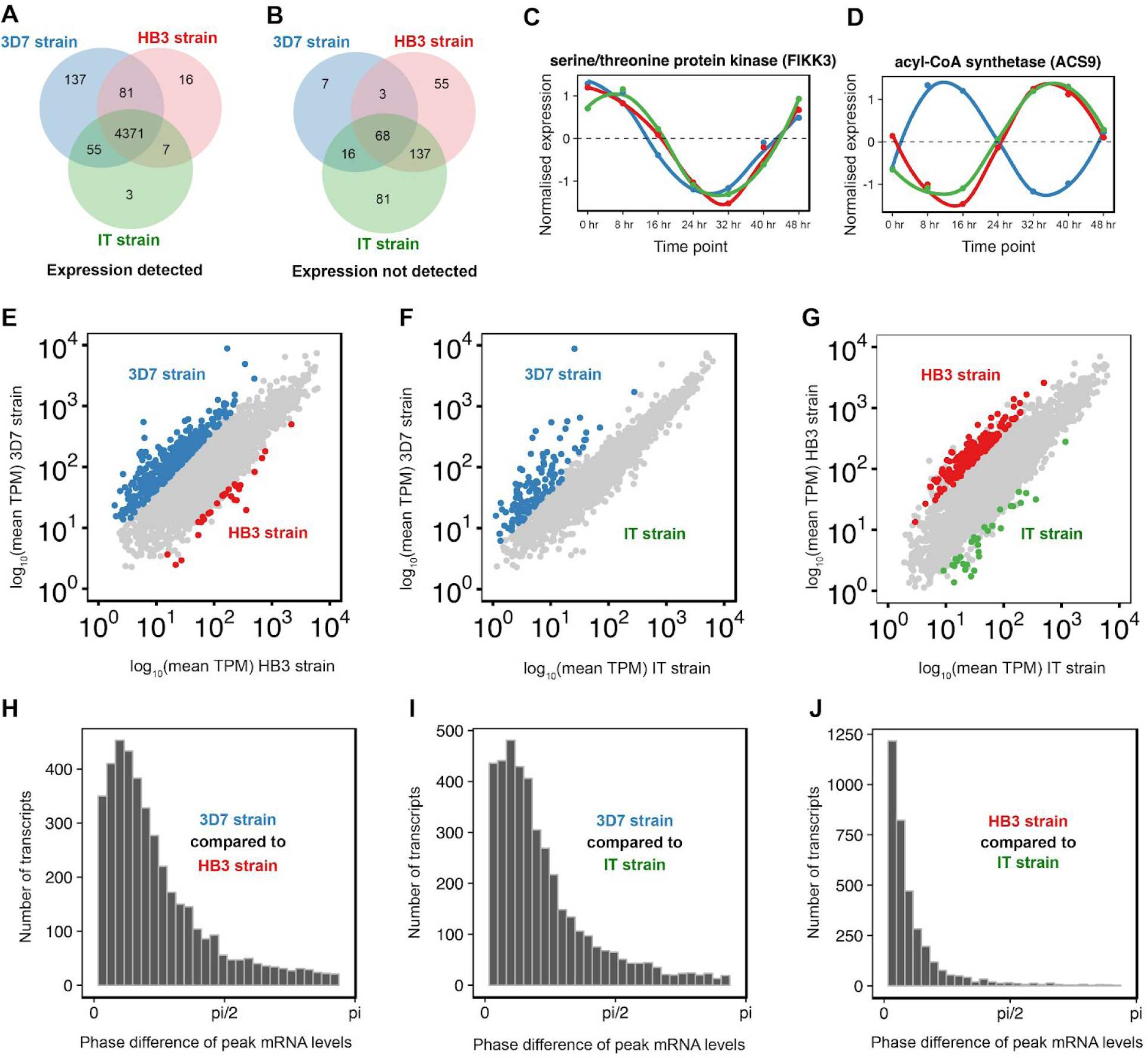
Comparison of blood stage transcriptomes from the 3D7, HB3, and IT strains of *P. falciparum*. A) Number of transcripts with expression detected in each of the three *P. falciparum* strains. Detection was based on a minimum expression of 5 TPM for at least one time point within the time course. Most transcripts (78%) were detected in all three strains. B) Number of transcripts with no expression detected in each of the three *P. falciparum* strains. Detection was based on a minimum expression of 5 TPM for at least one time point within the time course. C) Expression profile of the FIKK3 gene, which show similar timing/phase of expression in each of the three strains. D) Expression profile of the ACS9 gene, which shows a different timing/phase of expression in the 3D7 strain compared to both HB3 and IT. E) Identification of genes that are differentially expressed between the 3D7 strain and the HB3 strain. Coloured points highlight genes with at least a 2 fold difference in expression and FDR <= 0.05. F) Identification of genes that are differentially expressed between the 3D7 strain and the IT strain. G) Identification of genes that are differentially expressed between the HB3 strain and the IT strain. H) Phase (timing) differences in genes between the 3D7 strain and the HB3 strain. The mean phase difference was 0.747 radians (5.7 hours). I) Phase differences between the 3D7 strain and the IT strain. The mean phase difference was 0.697 radians (5.3 hours). J) Phase differences between the HB3 strain and the IT strain. The mean phase difference was 0.304 radians (2.3 hours).

From the DAFT-seq data we calculated differential abundance of mRNA between the three strains (**Table S24**, **Figure 6B**). Analysis of the functions associated with differentially expressed genes using GO terms showed that genes involved in the regulation of transcription show higher expression in 3D7 than HB3. Indeed, 15 out of 29 annotated ApiAP2 transcription factors were found to be detected at higher levels in 3D7 than in HB3 **(Figure S22, Figure S23)**. In addition, we only detect transcription of AP2-G, (*Pf3D7_1222600*), the key regulator of gametocytogenesis (59, 60), in 3D7, suggesting that the HB3 and IT strains used in this study cannot produce gametocytes **(Figure S25)**.

An example of a gene with similar expression in all 3 strains (FIKK3, *Pf3D7_0301200*) is shown in **Figure 6C**, while **Figure 6D** shows differing expression profiles for ACS9 (putative acetyl-CoA synthetase, *Pf3D7_062780*), which is a member of an expanded gene family in *P. falciparum* with variable gene expression depending on the cell line. As reported previously (Llinás et al NAR 2005), the timing of gene expression was highly correlated between *P. falciparum* strains (**Figure 6E-G**), with few genes showing large differences in expression timing (**Figure 6H-J**). HB3 and IT showed the strongest correlation in timing of gene expression, whereas 3D7 and IT were more correlated in amplitude (**Figure S24**, **Table S25**).

Finally, we identified UTRs and spliced transcripts for all three strains. Our UTR-detection pipeline identified fewer UTRs (∼3,800) for HB3 and IT than for 3D7 **(Tables S26, S27; Figure S25)**, which is likely to be due to the higher quality and completeness of the underlying 3D7 reference genome sequence. Where gene expression was detectable in more than one strain, the length of the UTR calls were largely comparable (R ∼ 0.8). There were slightly more splice sites observed in HB3 (9,231) than in the other two strains (8,688 in 3D7, 7,627 in IT; **Tables S28, S29**), and we also detected more exitrons in HB3 (248) and IT (197) (**Table S30, S31**) than in 3D7 (155). It is not yet clear why there is this variation between the strains. We also observed that the relationship found for 3D7 between expression of head-to-head and tail-to-tail genes was similar across all three strains (**Figure S26, Tables S32-S35**).

## Discussion

The combined use of DAFT-seq, 5UTR-seq and PacBio cDNA protocols enabled the *P. falciparum* transcriptome to be systematically re-evaluated without the GC-content biases induced by PCR amplification and random priming. These new data revealed that 89% of the 3D7 genome is transcribed throughout the IDC. The 5’ and 3’ UTRs we identified (averaging roughly 600 nt each) are large compared to the average length of a protein-coding sequence (around 2,000 nt), suggesting that much of the genome is not intergenic “space”, but is associated with some function or biochemical activity. A key feature of the DAFT-seq data is that the coverage is near continuous from the extreme 5’ end of a transcript through to the 3’ end, allowing linkage of features throughout a transcript. We suggest that this unambiguous linking of the 5’ and 3’ UTRs to the main gene body makes our set of predicted 5’ and 3’ UTRs highly useful to researchers wanting to identify mRNA boundaries.

These newly defined transcription start sites will be especially useful to researchers wanting to generate knock-out or tagged parasites lines, and to researchers using single cell RNA-seq protocols that target either the extreme 5’ or 3’ ends of mRNAs. In other eukaryotic systems both the sequence and structure of the 5’ and 3’ UTRs of transcripts are known to play key roles in gene regulation (61, 62). The relative position and sequence of the UTRs in *Plasmodium* parasites is therefore likely to be constrained by functional requirements, in which particular nucleotides will be crucial for the formation of tertiary structures in the mRNA molecules recognised by RNA-binding proteins. Perhaps structural constraints imposed by tertiary structures of 3’ UTRs affect the amino acid usage in overlapping protein-coding sequences? We found relatively few differences between the 5’ and 3’ UTR sequences used by the three strains of *P. falciparum* investigated in this study, suggesting that UTR position or sequence motifs are highly conserved between different parasites.

The extensive and internally validated mapping of TSSs in these data allowed us to extend this knowledge and conclude that there is a “zone” of transcriptional initiation, within which there can be temporal variation in TSS position throughout the IDC. Combining these new data with additional genome-wide data sets such as nucleosome occupancy (15) and ATAC-seq (16, 63) may lead to further mechanistic insights into the control of transcription initiation in *P. falciparum*. Perhaps different regions of DNA are accessible or bound by different regulatory proteins during the IDC, or certain regions of the UTRs are important for the binding of post-transcriptional regulators that could stabilise or destabilise the transcript.

The genome-wide locations of 5UTR-seq TSSs we determined differ significantly with those in another recent study (43) which reported that 49% of all “TSSs blocks” were downstream of the annotated ATG start codons. In contrast, 90% of TSSs (analysed using all data and a 2,000 nt upstream window) and 89-94% of the TCs (analysed per time point) we found mapped outside protein-coding sequences. We suggest that these differences may be due to methodological differences at two key points in the protocols used. Firstly, we selected polyA+ transcripts rather than those solely containing a 5’ cap. Secondly, Adjalley et al. use PCR amplification in their library production protocols; PCR amplification of AT-rich DNA usually enriches for GC-rich molecules and depletes AT-rich ones (64, 65). This bias would be likely to enrich for TSS reads in relatively GC-rich exons (∼80% AT) compared to the most AT-rich non-coding regions (∼90% AT), where continuous UTR sequences were only reliably detected once we omitted the PCR amplification step from our protocol.

The strand-specific nature of the DAFT-seq protocol allows unequivocal confirmation of transcription from the non-coding strand of genes. However, the more precisely defined boundaries of transcriptional units suggests that many of the “antisense transcripts” observed in previous studies are either TSSa-RNAs or overlapping 3’ UTRs. There are at least two biological explanations for the presence of TSSa-RNAs. One explanation is that they are a normal byproduct of promoter activity, similar to that described previously for higher eukaryotes (57). A second explanation is that these transcripts have some function in gene regulation, which we speculate could include binding by regulatory proteins containing an RNA recognition motif (RRM), many of which are encoded in the *P. falciparum* genome (66), but which are largely uncharacterised (67). Additionally, our identification of independently regulated ncRNAs may help future studies of gene regulation, as seen with an ncRNA antisense to GDV1 (68) (which can be observed in our data, **Figure S27**), which is involved in the regulation of sexual commitment.

The putative exitrons detected suggest an additional mechanism of regulatory complexity, enabling the parasite to generate multiple protein isoforms from even single exon genes. This is of particular relevance for genes that have multiple binding partners or domains, such as transcription factors. Even for proteins with single interactors, inclusion or exclusion of key protein sequence may determine the activity or localisation of the protein.

Together, the complementary approaches applied in this study allow a refined description of the transcriptional landscape of *P. falciparum* and demonstrate that very little of the densely packed *P. falciparum* genome can be considered inactive or redundant. These precise definitions of 5’ and 3’ UTRs will be useful in guiding the definition of regions for amplification in experimental gene expression studies. In addition, the analysis identifies multiple regions where transcriptional units are very close or even overlap. This information will be of particular use for experimental genetics approaches aimed at deleting or altering specific genes, by highlighting regions where the insertion of gene modification constructs may have unintended functional consequences on adjacent genes. We anticipate that identifications of these mRNA transcript overlaps will motivate reanalysis of unexplained mutant phenotypes, where extended gene models can provide additional insights into the regulation of adjacent genes.

## Data Availability

All sequencing data generated and analyzed in this work have been deposited in the European Nucleotide Archive (ENA) for public access under study accession number is ERP001570. The processed data for transcription start sites are also being made publicly available through the Plasmodium genome resource PlasmoDB.org (69).

## Acknowledgements

We would like to acknowledge Chris Newbold for helpful discussions throughout this project. We would like to acknowledge Ulrike Böhme for help regarding genome annotation and Mandy Sanders for coordinating the sequencing. We would like to acknowledge the Sanger DNA Pipelines Bespoke Library team for quantifying and loading of sequencing libraries. We would like to thank Adam Reid, Martin Hunt, Richard Bartfai and David Conway for helpful advice.

