## Supplementary tables for "Refining the transcriptome of the human malaria parasite *Plasmodium falciparum* using amplification-free RNA-seq"

**This PDF file includes:**

Figures S1-S27
Supplementary text
References

### Supplementary figures

Figure S1: DAFT-seq schematic
Figure S2: PolyA+ vs Random hexamer priming of mRNA in the DAFT-seq protocol
Figure S3: Method for calling UTRs from DAFT-seq coverage data
Figure S4: Comparison of Chappell et al. 5’ UTRs with the Caro et al. study
Figure S5: Comparison of TSSs with the Adjalley et al. study
Figure S6: 5UTR-seq schematic
Figure S7: Method for calling 5’ UTRs with 5UTR-seq data
Figure S8: Alternative 5’ UTR sets
Figure S9: Homopolymer tracts
Figure S10: PacBio reads show the same features of the transcriptome as DAFT-seq data
Figure S11: Distinct temporal changes of TSS usage in the IDC (KAHRP)
Figure S12: The TSS frequency and shape landscape for 3D7
Figure S13: TSS motifs in introns and exons
Figure S14: Sequence features of sharp and broad promoters
Figure S15: ApiAP2 motif occurrences relative to predicted TSSs and annotated translation start sites in the 3D7 strain
Figure S16: ApiAP2 motifs around the KAHRP gene
Figure S17: Exitrons in PacBio reads
Figure S18: Alternative splicing in PacBio reads
Figure S19: TSS associated non-coding RNAs
Figure S20: Long non-coding RNAs
Figure S21: Alignment of DAFT-seq data to microarray data
Figure S22: ApiAP2 expression in the 3 *P. falciparum* strains
Figure S23: AP2-G expression in the 3 *P. falciparum* strains
Figure S24: Amplitude change distributions of all genes in the 3 *P. falciparum* strains
Figure S25: Coverage-based UTRs for the 3 *P. falciparum* strains
Figure S26: Analysis of expression patterns of adjacent gene pairs for the 3 *P. falciparum* strains
Figure S27: A spliced non-coding RNA opposite GDV1

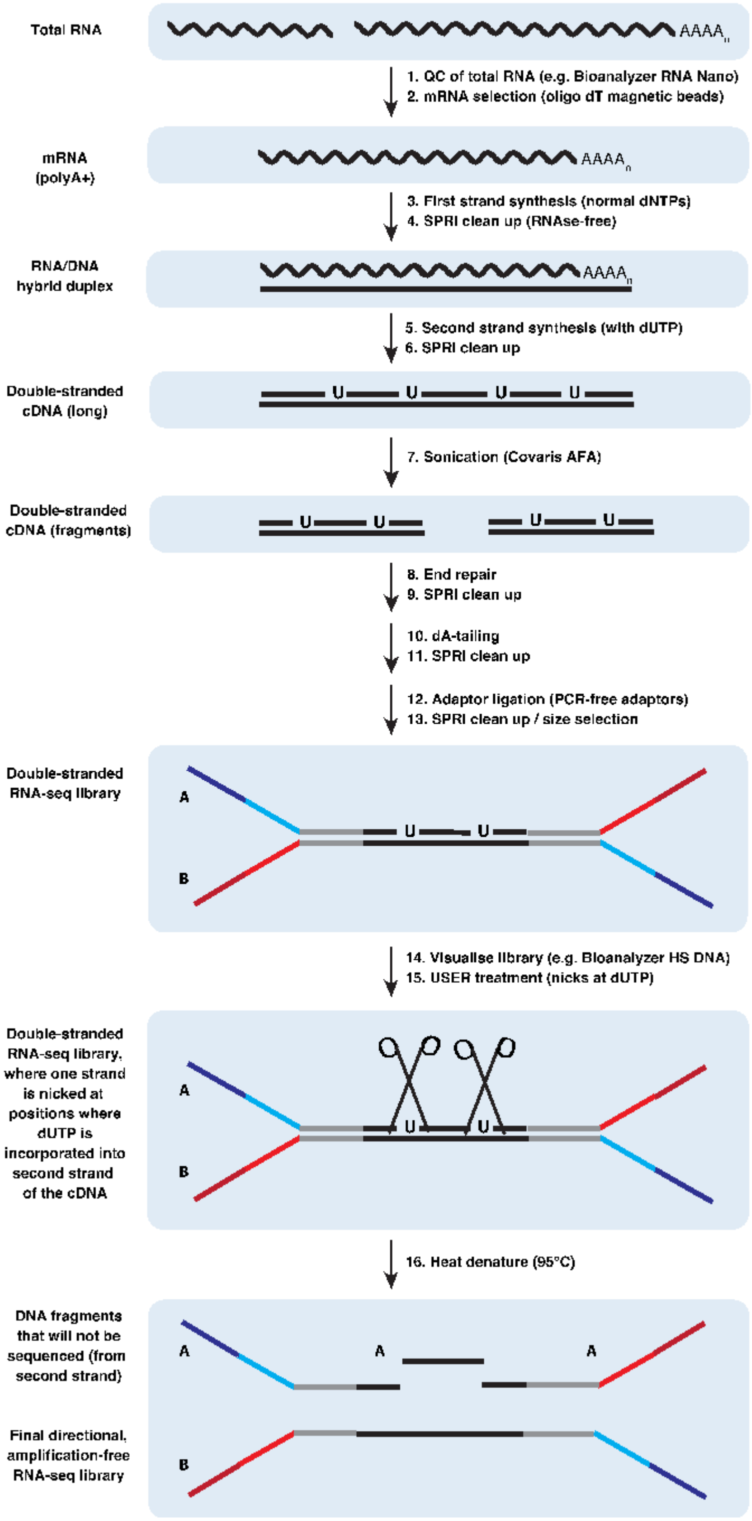


**Figure S1: DAFT-seq schematic**

**Figure S1: DAFT-seq schematic**

Overview of the directional, amplification-free transcriptome sequencing (DAFT-seq) protocol. The input to the protocol is quality-controlled total RNA; poly A+ RNA (mostly mRNA) is then isolated using magnetic oligo (d)T beads. Reverse transcription of full-length mRNA is initiated using oligo d(T) primers, and is followed by a cleanup of the reaction mix using SPRI beads. Next, second strand cDNA is synthesised; dUTP is included instead of dTTP in the dNTP mix to enable retention of directional information. Standard end repair and dA tailing reactions are then performed. Adaptor ligation was modified to include longer "PCR-free" adaptors (similar to those in Kozarewa *et al*. (2009)), which include all the sequence required for Illumina sequencing, allowing the PCR step to be omitted. After SPRI cleanup the second strand cDNA containing dUTP residues is digested using the USER enzyme mix, generating the final directional libraries.

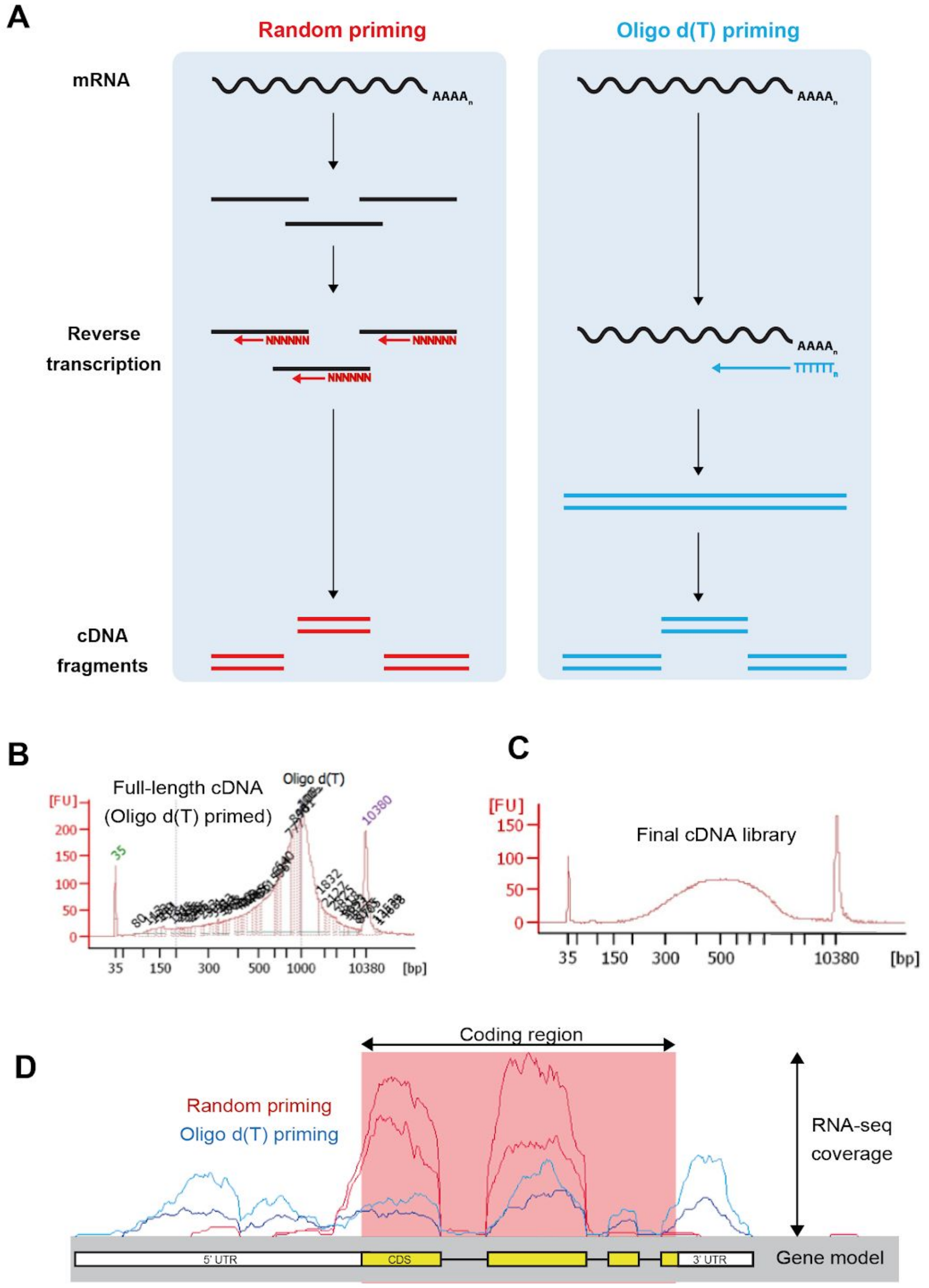


**Figure S2: PolyA+ vs Random hexamer priming of mRNA in the DAFT-seq protocol**

**Figure S2: PolyA+ vs Random hexamer priming of mRNA in the DAFT-seq protocol**

**A)** Left: cDNA synthesis using random priming of mRNA fragments. This is the approach used in most RNA-seq studies, where fragmentation of mRNA is used to improve the evenness of coverage along the length of the transcript. This approach produces uneven coverage in *P. falciparum* RNA-seq libraries, as the mRNA is not primed in AT-rich regions. Right: cDNA synthesis using oligo d(T) priming of full-length mRNA. This approach has previously been shown to have a 3' end bias in model organisms, but provides more even coverage across genes and in extremely AT-rich regions when used for directional PCR-free *P. falciparum* RNA-seq libraries, as priming only needs to take place once in the polyA+ tail rather than within every AT-rich region.

**B)** Bioanalyzer trace (on HS DNA chip) of full-length cDNA primed using oligo d(T) primers. The peak to the right is >1kb.

**C)** Bioanalyzer trace (on HS DNA chip) of final cDNA library from a DAFT-seq protocol, using cDNA primed with oligo d(T). The average size is appears around 500 bp, though the libraries appear larger than the "true" size, due to the longer "Y-shaped" tails on the PCR-free sequencing adaptors.

**D)** RNA-seq coverage traces of PCR-free cDNA libraries made from cDNA primed using random primers only (red trace) or with oligo d(T) primers (blue trace). The libraries were made using the same RNA, so differences in coverage are related to differences in cDNA priming. The random primer library (red trace) has most of its coverage concentrated in the relatively GC-rich coding region of this gene (red box, average AT content of protein-coding sequence is ~80%), with little coverage outside this region (average AT-content of non-coding regions is >90%). Splice sites are seen as sharp drops in coverage within the coding region. The oligo d(T)-primed library shows a flatter coverage distribution across the coding region and adjacent non-coding regions, which are annotated as 5' and 3' UTRs. This more even coverage across genomic sequences variable and extreme AT-content enables annotation of UTRs on a genome-wide scale.

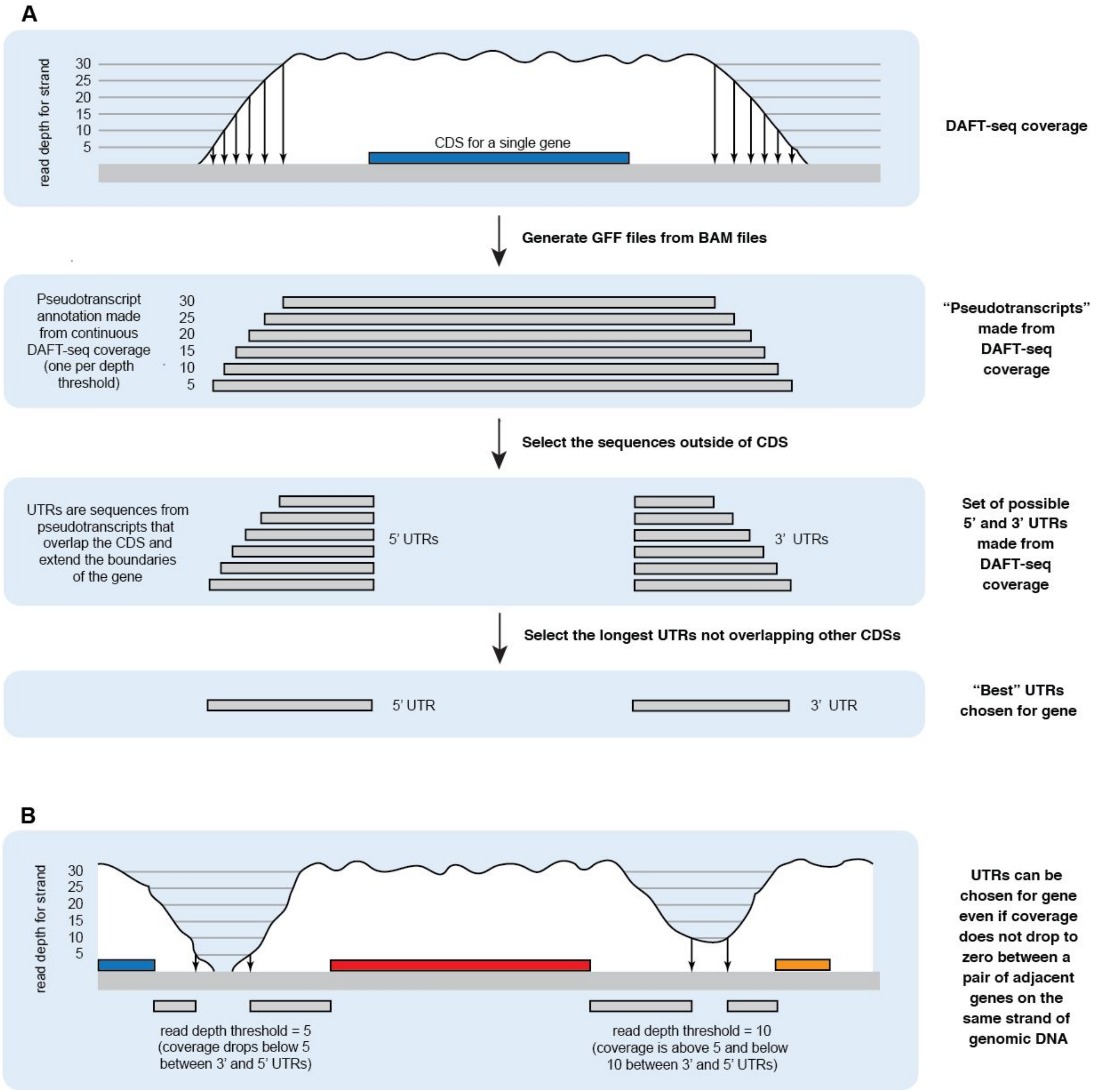


**Figure S3: Method for calling UTRs from DAFT-seq coverage data**

Schematic view of the method for calling 5' and 3' UTRs based only on DAFT-seq coverage.

**A)** Blocks of continuous DAFT-seq data overlapping with each gene model were used to make "pseudotranscripts" at a range of different coverage thresholds (5 to 100 reads depth, in intervals of 5). The coding sequence is "cut out" to generate the 5' and 3' UTRs.

**B)** In cases where adjacent UTRs on the same strand overlap, a more stringent coverage threshold is used to distinguish the two UTRs.

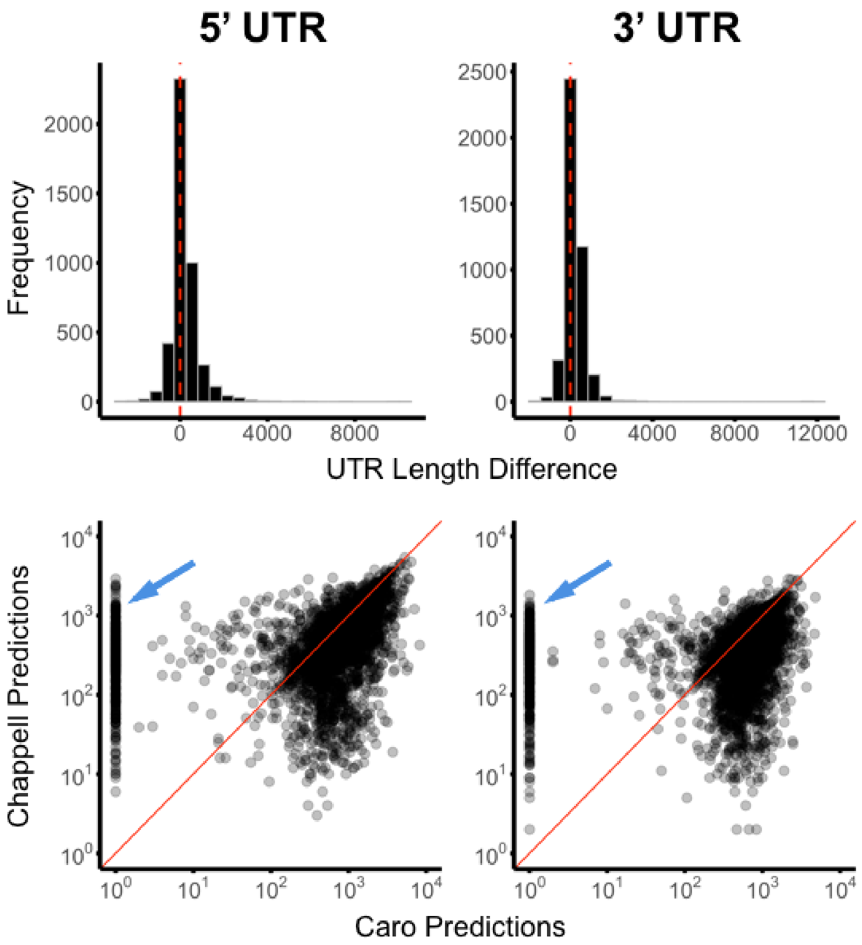


**Figure S4: Comparison of UTRs with the Caro *et al.* study**

5' and 3' UTR predictions from Caro *et al.* were downloaded and compared to 5' and 3' UTR predictions in this study. The top row displays the difference in lengths for the whole set, while the bottom row directly compares lengths between individual genes. The longest 5' and 3' UTR prediction for each gene from Caro *et al.* were compared to our predictions. Caro *et al.* predictions were on average 230 and 208 bp longer while Pearson correlation estimates were 0.59 (95% CI 0.57,0.61) and 0.36 (95% CI 0.33,0.38) for 5' and 3' UTR comparisons, respectively. Notably, despite on average longer UTR lengths, 969 5' UTRs and 940 3' UTRs (indicated by the blue arrows) from Caro *et al.* were predicted to have lengths of zero (i.e. transcription begins at the translation start site), but had non-zero length UTRs in our data set. The slope of the solid red line is equal to one.

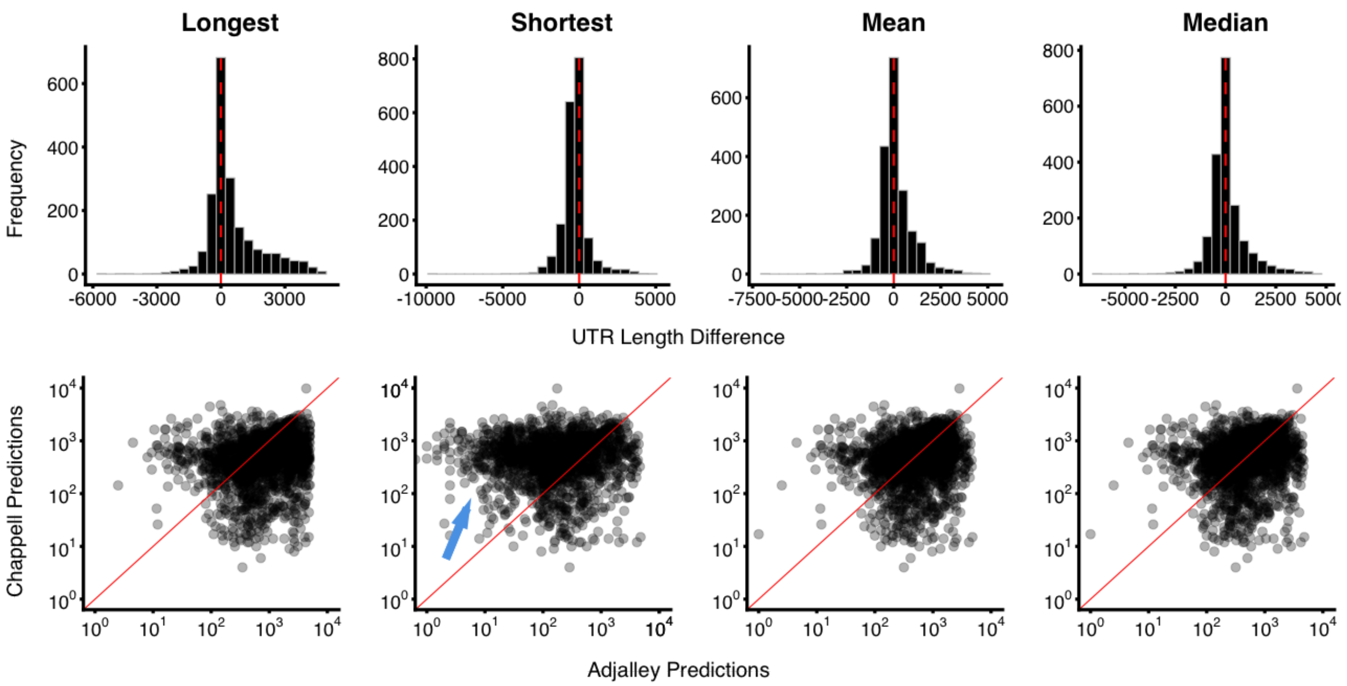


**Figure S5: Comparison of TSSs with the Adjalley *et al.* study**

TSS predictions from Adjalley *et al.* were compared to TSS predictions in this study. Longest, shortest, mean, and median 5’ UTR length predictions for each gene were calculated from TSS predictions in Adjalley *et al.* 5’ UTRs annotated for the same gene were directly compared to 5’ UTR predictions in this study. Predictions were, on average, 562, -263, 117, and 84 bp longer for calculated longest, shortest, mean, and median 5’ UTRs, respectively (minus sign indicates that Chappell *et al.* predictions were, on average, longer). Pearson correlation estimates were 0.05 (95% CI 0-0.9), 0.15 (95% CI 0.10-0.19), 0.14 (95% CI 0.10-0.19), and 0.15 (95% CI 0.10-0.19) for calculated longest, shortest, mean, and median 5’ UTRs respectively. The blue arrow indicates an enrichment of shorter 5’ UTR predictions in the Adjalley *et al.* dataset.

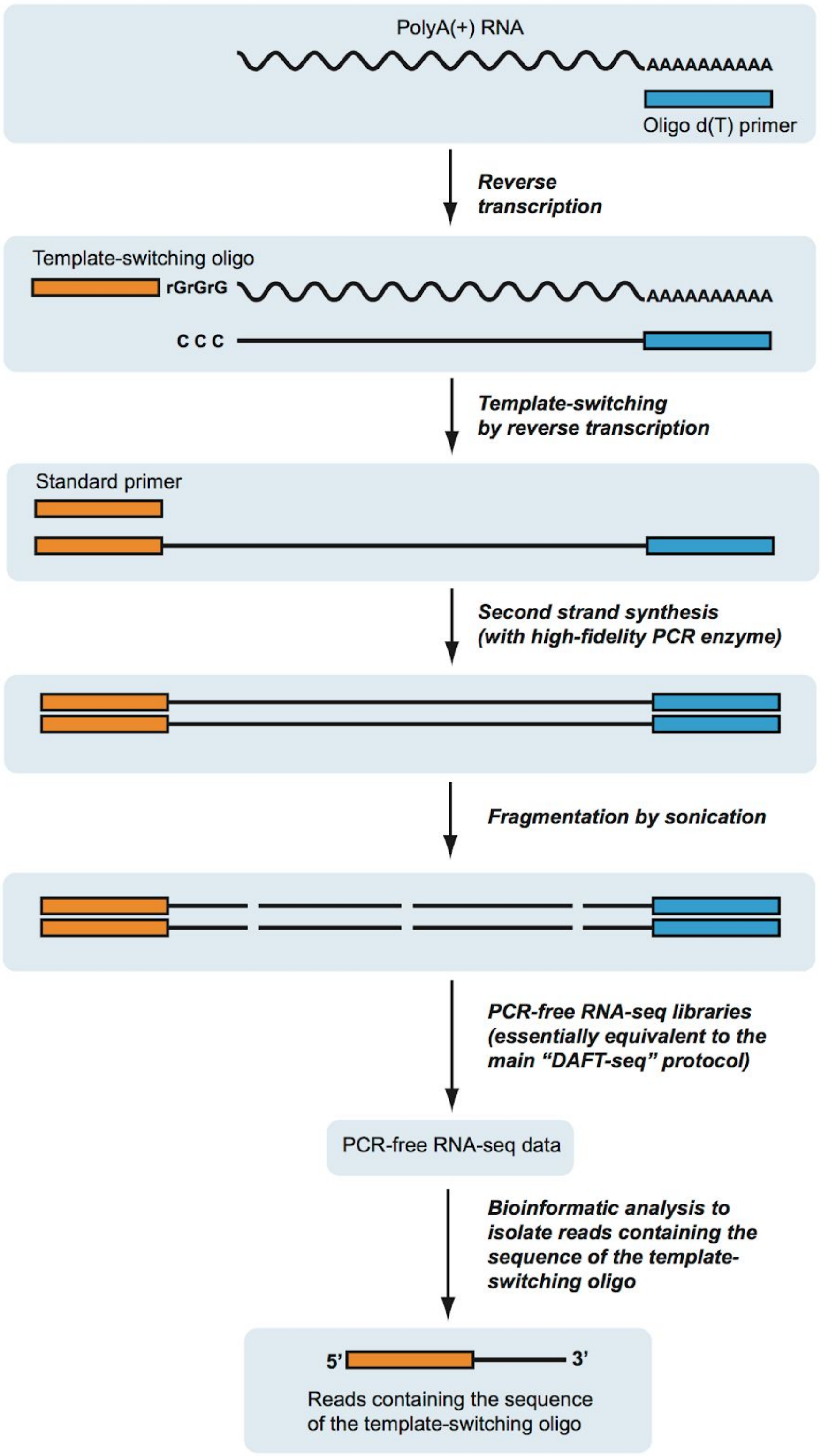


**Figure S6: 5UTR-seq schematic**

The 5UTR-seq protocol uses similar molecular biology to the DAFT-seq protocol to generate full-length cDNA from polyA+ mRNA molecules, but additionally includes a template-switching oligo (TSO). A number of modified mouse leukemia virus (MMLV) reverse transcriptases can add non templated Cs to the 3' end of first strand cDNA where a cap is present at the 5' end of the mRNA. In the presence of the TSO the reverse transcriptase switches template sequence, the effect of which is to add a common oligo handle to the end of all full-length cDNAs. This TSO "tags" the position in the cDNA that corresponds to the extreme 5' end of the original mRNA, and can be used to infer the position of the TSS.

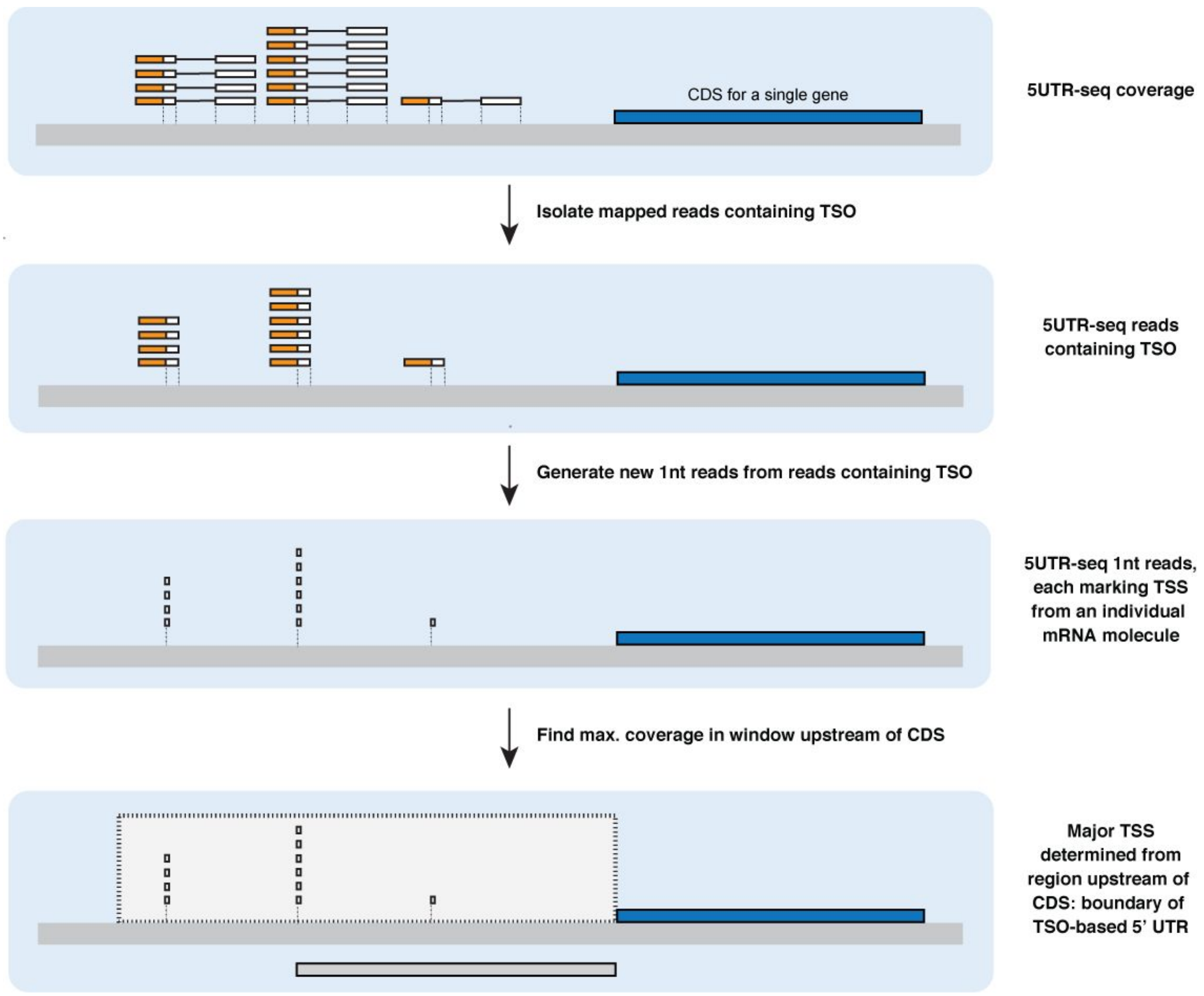


**Figure S7: Method for calling 5' UTRs with 5UTR-seq data**

5UTR-seq reads are mapped to the genome using paired reads, using a mapper (SMALT) that enables "soft-clipping" (overhang) of the TSO, which will not map to the genome. The mapped reads containing the TSO are then isolated (using BLAST), and are processed further to generate single nt coordinates that correspond to the first transcribed bases. For each gene, the reads corresponding to the TSSs of individual molecules are analysed for a defined window upstream of the coding sequence. The relative heights of stacks of reads correspond to relative frequencies of TSS usage.

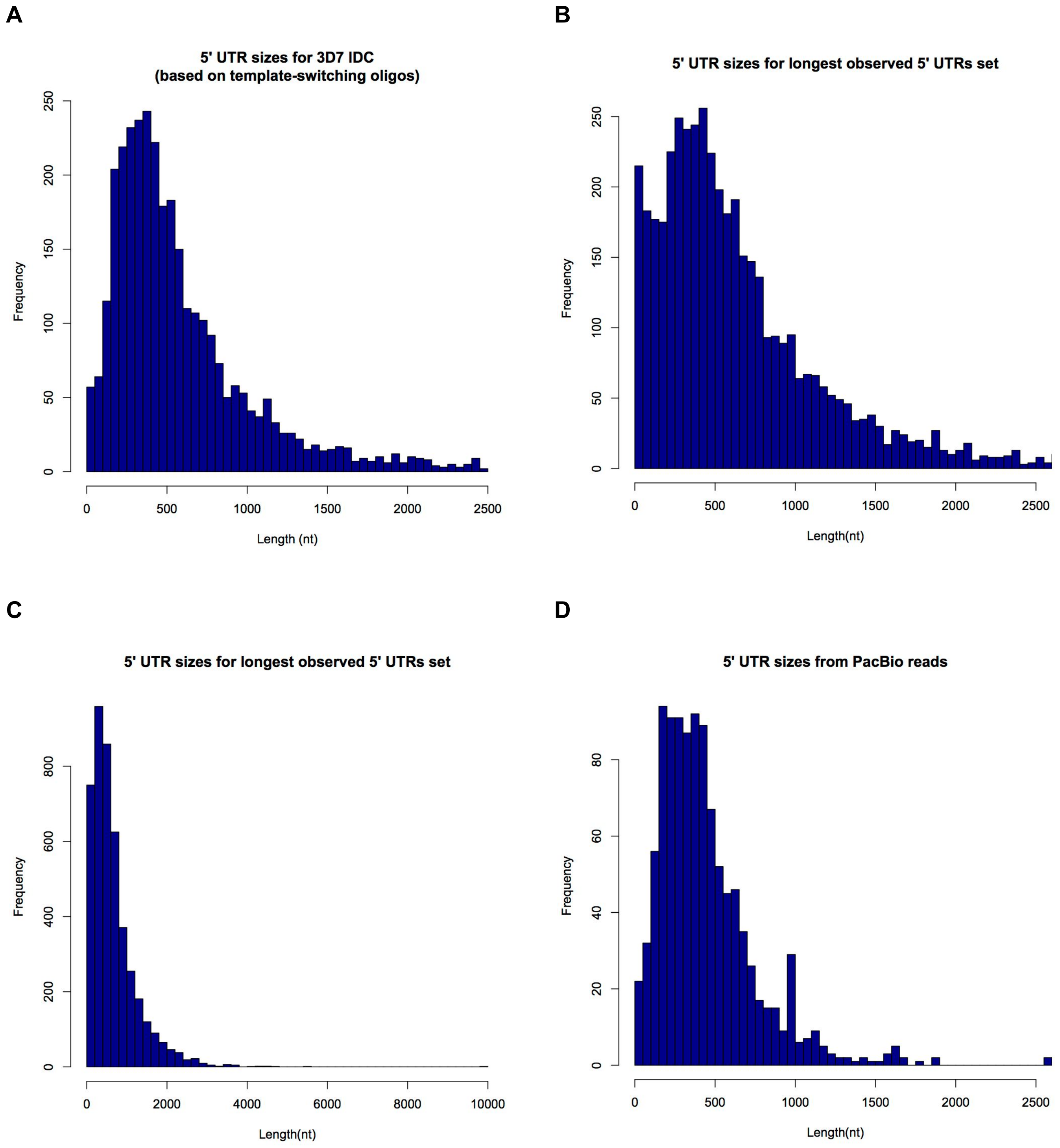


**Figure S8: Alternative 5’ UTR sets**

**A)** TSO-based 5’ UTRs (maximum length set to 2,500 nt in detection)
**B)** Longest observed 5’ UTRs from combined set (shown to 2,500 nt)
**C)** Longest observed 5’ UTRs from combined set (shown to 10,000 nt)
**D)** PacBio TSO-based 5’ UTRs (maximum length set to 2,500 nt in detection)

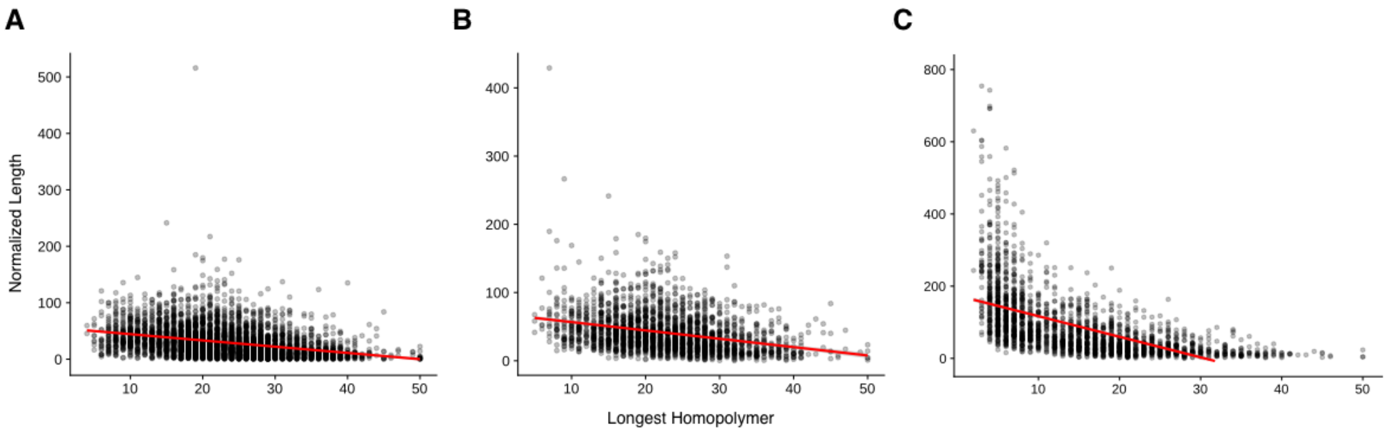


**Figure S9: Homopolymer tracts upstream of predicted 5' UTRs**

**A)** The 5' UTR length of all 5' UTR sequences plus 100 bp ustream, normalized by the longest homopolymer tract plotted as a function of the longest homopolymer tract found within. There is a modest, but significant negative correlation of -0.320 (p-value of 4.10e-56; 95% confidence intervals of -0.357 to -0.283) suggesting that where there are longer homopolymer tracts, the predicted 5' UTRs are also shorter.

**B)** The 5' UTR length of 5' UTR sequences that were "repaired" plus 100 bps ustream, normalized similar as in A. Again, we see a modest, but significant negative correlation of -0.335 (p-value of 2.33e-115; 95% confidence intervals of -0.361 to -0.308).

**C)** 100 bp ustream of 5' UTR sequences that were repaired, normalized similar as in A. We see an even more significant negative correlation of -0.561 (p-value of 5.26e-191; 95% confidence intervals of -0.588 to -0.532). We interpret these results as showing that low complexity regions within *P. falciparum* intergenic regions are responsible for causing gaps in coverage that may bias UTR predictions towards shorter lengths.

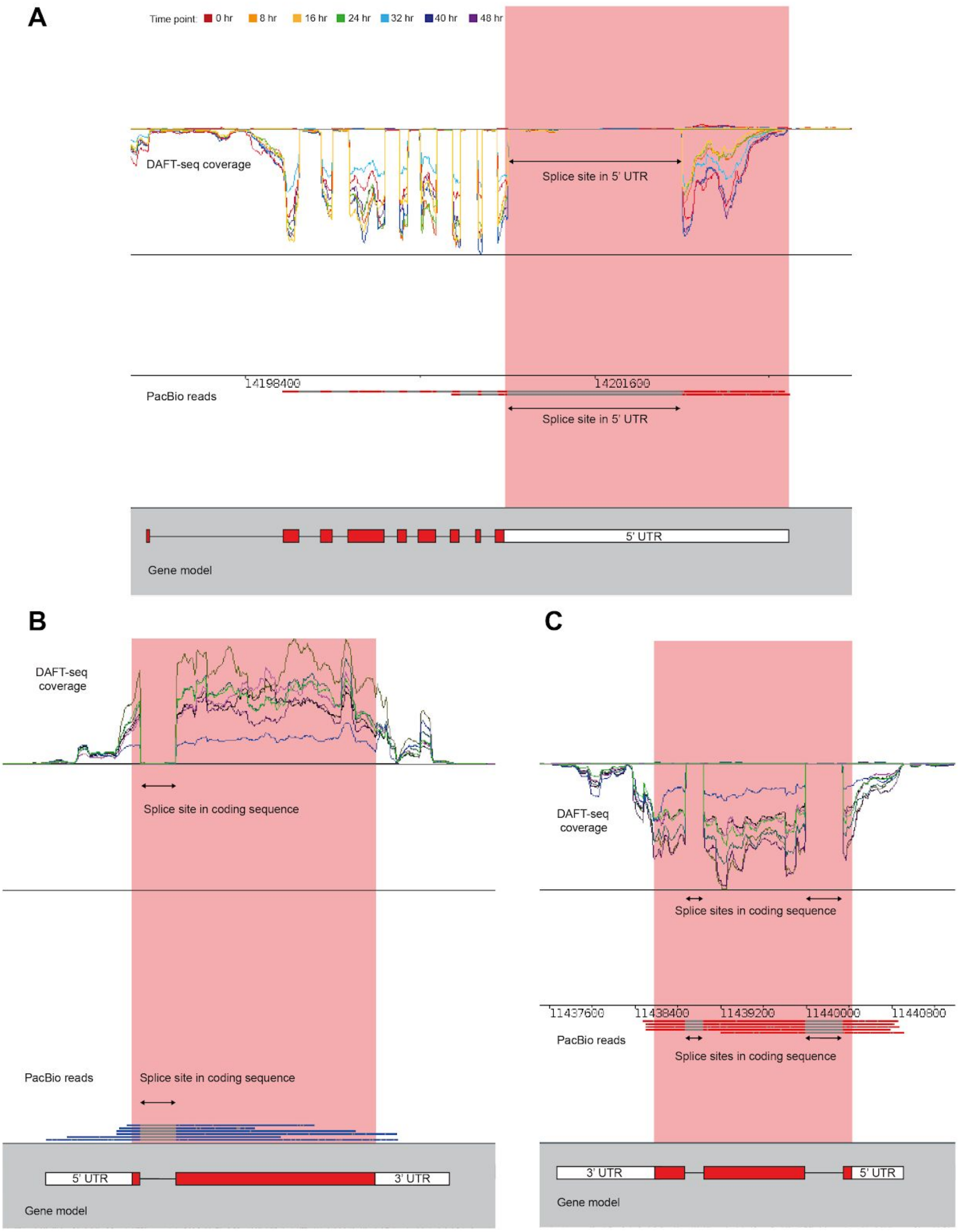


**Figure S10: PacBio reads show the same features of the transcriptome as DAFT-seq data**

**A)** The longest 5 'UTR (Pf3D7_1136500) in the PacBio data set (middle panel) is very similar to the 5' UTR of the same gene in the DAFT-seq data set (top panel)
**B)** The splice sites found in the PacBio reads for the gene Pf3D7_0917900 (middle panel) are the same as those found using the DAFT-seq data set (top panel)
**C)** The isoform structure found in the PacBio reads for the gene Pf3D7_1008700 (middle panel) is the same as that found using the DAFT-seq data set (top panel)

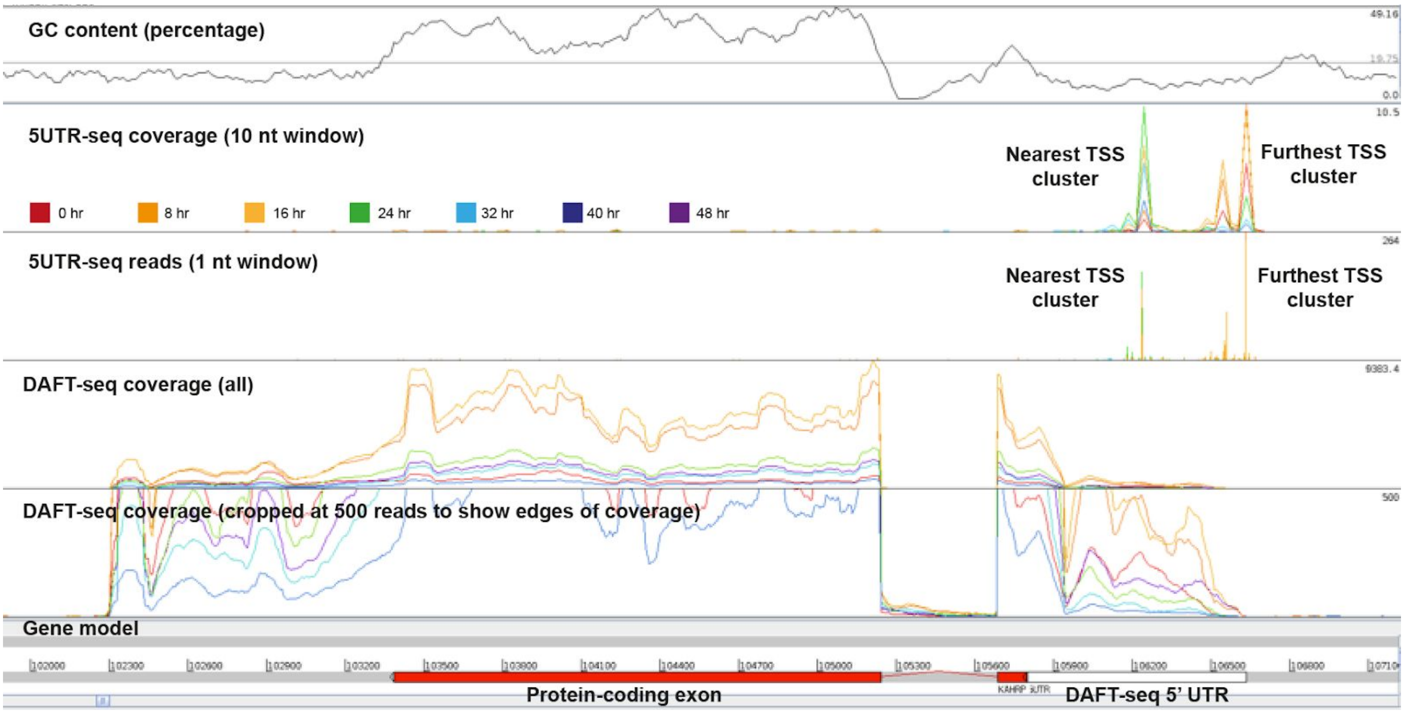


**Figure S11: Distinct temporal changes of TSS usage in the IDC (KAHRP)**

There is a change in the relative usage of TSSs between the clusters of TSSs that are nearest and furthest from the start codon of the KAHRP gene between different time points in the IDC (see key for colours of different time points).

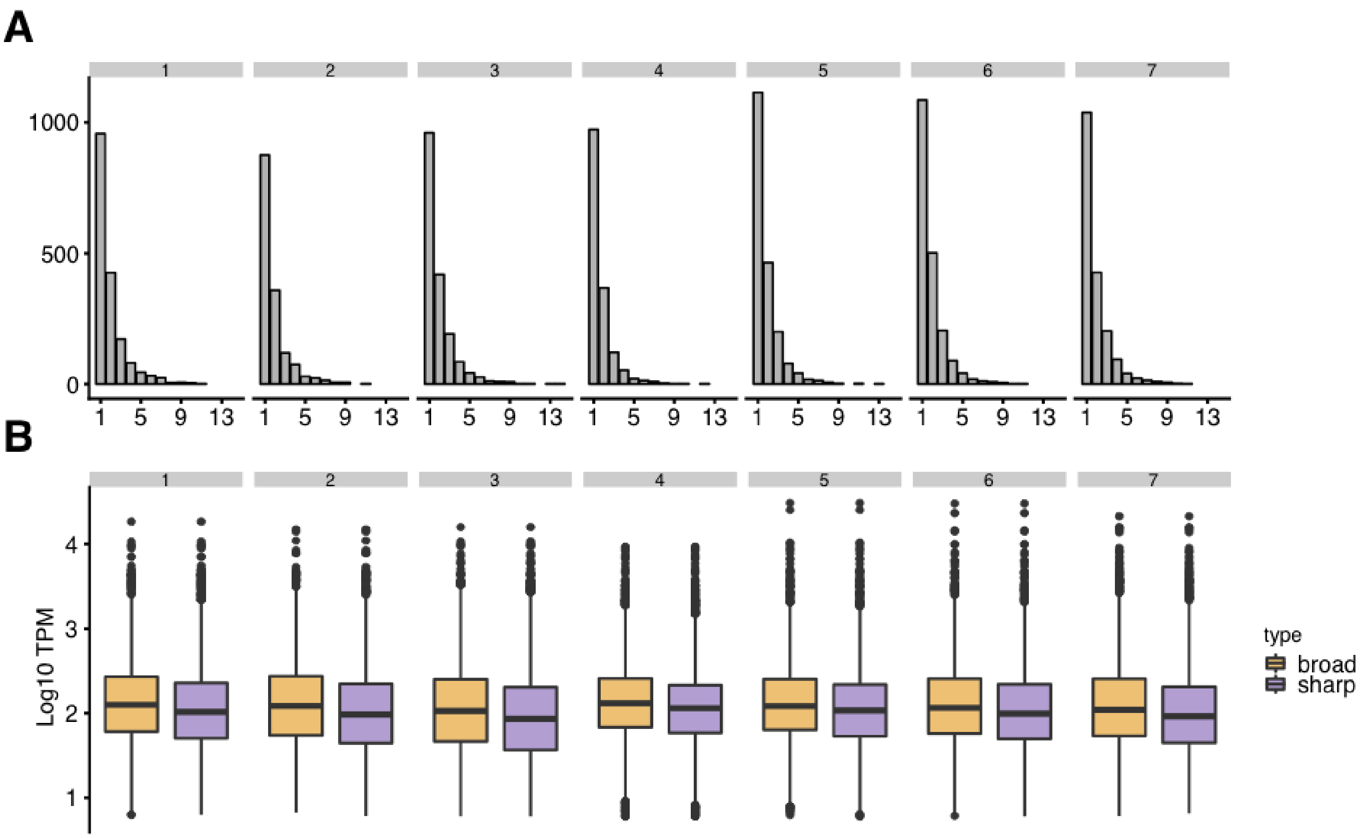


**Figure S12: The TSS frequency and shape landscape for 3D7**

**A)** The number of annotated tag clusters (TCs) per gene, per time point (7 panels). Most genes are annotated with just a single TC for each time point (tallest bar,~1000 genes in each time point). However, many genes also have multiple annotated TCs per time point.

**B)** Expression of genes annotated with broad promoters are significantly (Welch Two Sample t-test) and consistently more highly expressed than genes annotated by sharp promoters (p-values <1e-8 for all time points).

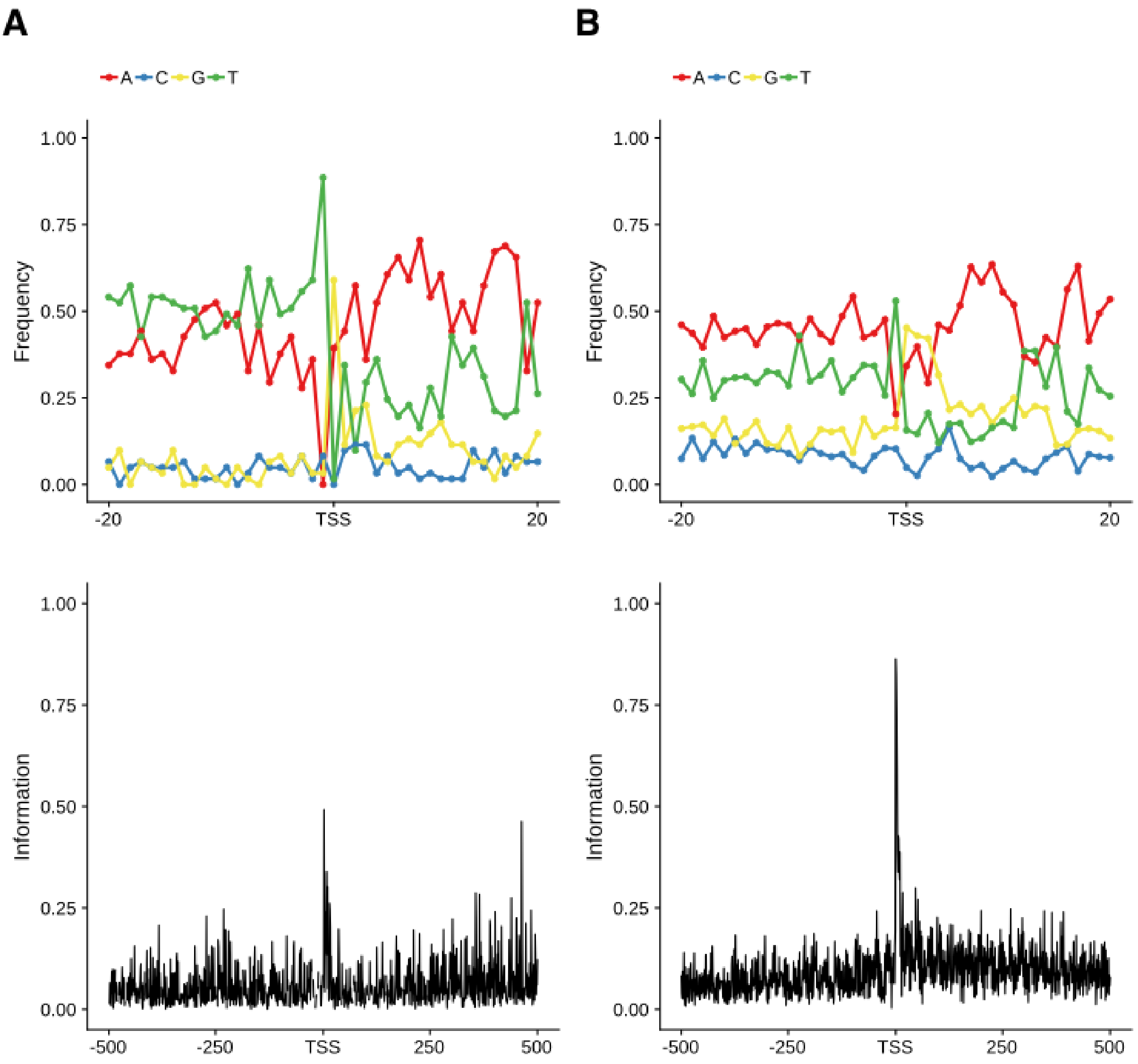


**Figure S13: TSS motifs in introns and exons**

**A)** Top panel: average nucleotide frequency for each position surrounding predicted TSSs found within introns (n=61). Lower panel: calculated information content for each position.

**B)** Top panel: average nucleotide frequency for each position surrounding predicted TSSs found within exons (n=389). Lower panel: calculated information content for each position.

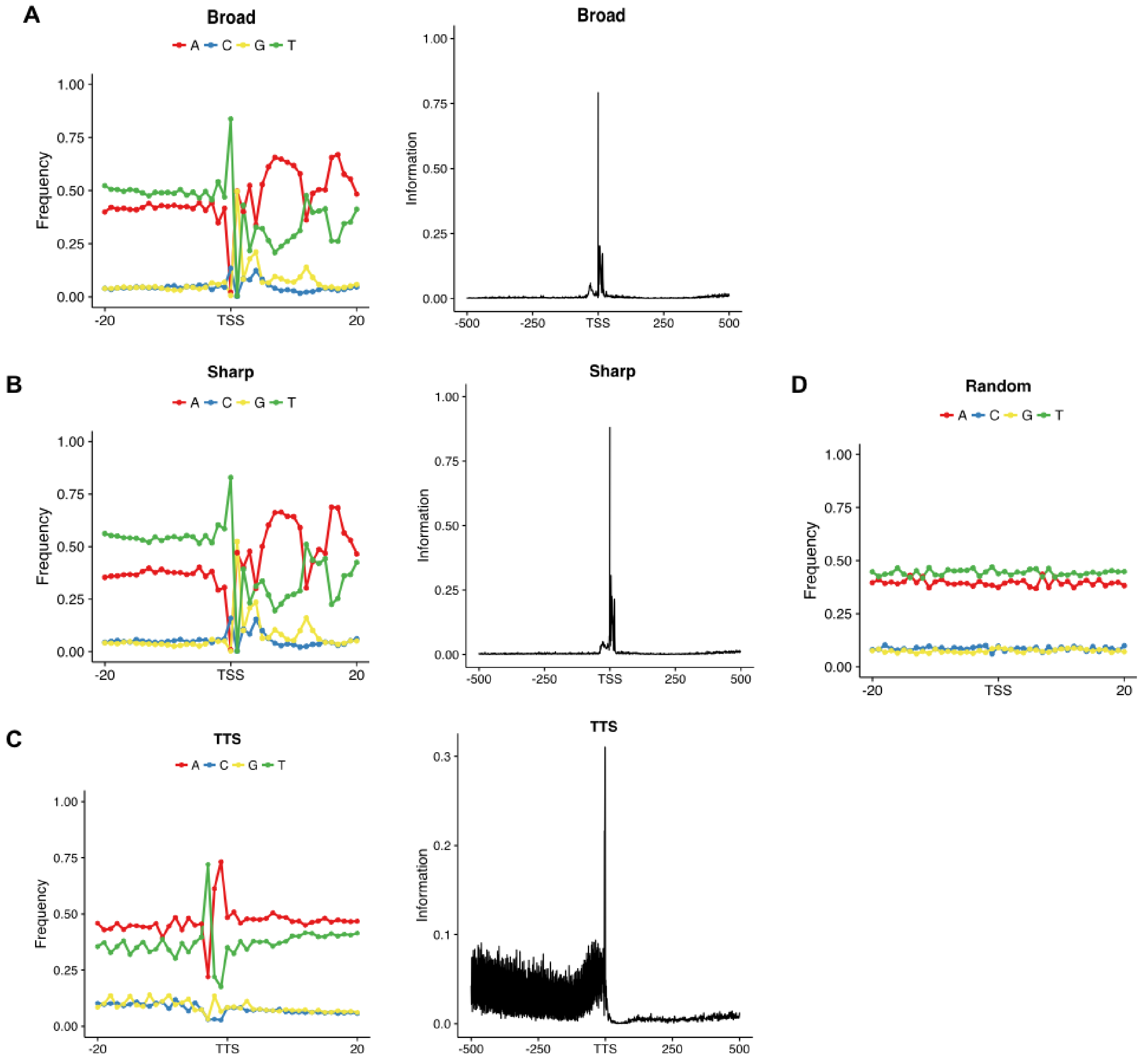


**Figure S14: Sequence features of sharp and broad promoters**

**Figure S14: Sequence features of sharp and broad promoters**

**A)** Left panel: nucleotide frequency surrounding TSSs found in broad promoters, in a 40 bp window centred on the TSS. Broad promoters show distinct sequence patterns surrounding TSSs. A consensus “TG/AG/A” appears surrounding the -1 and +1 nucleotides. Right panel: information content for each position within a 1000 bp window. We observe a peak at defined TSSs and a much smaller peak within 50 bp upstream.

**B)** Left panel: nucleotide frequency surrounding TSSs found in sharp promoters, in a 40 bp window centred on the TSS. Sharp promoters show nearly the same sequence patterns as broad along with a similar consensus sequence.content for each position within a 1000 bp window. Right panel: information content for each position within a 1000 bp window. As with broad promoters, this shows a peak at defined TSSs and a much smaller peak within 50 bp upstream.

**C)** Left panel: nucleotide frequency surrounding transcription termination sites (TTSs). TTSs have subtle, but distinguishable sequence patterns.Right panel: information content for each position within a 1000 bp window, which shows a major change in variability following the termination site.

**D)** Randomly chosen sites throughout the genome show a uniform distribution of nucleotide frequencies.

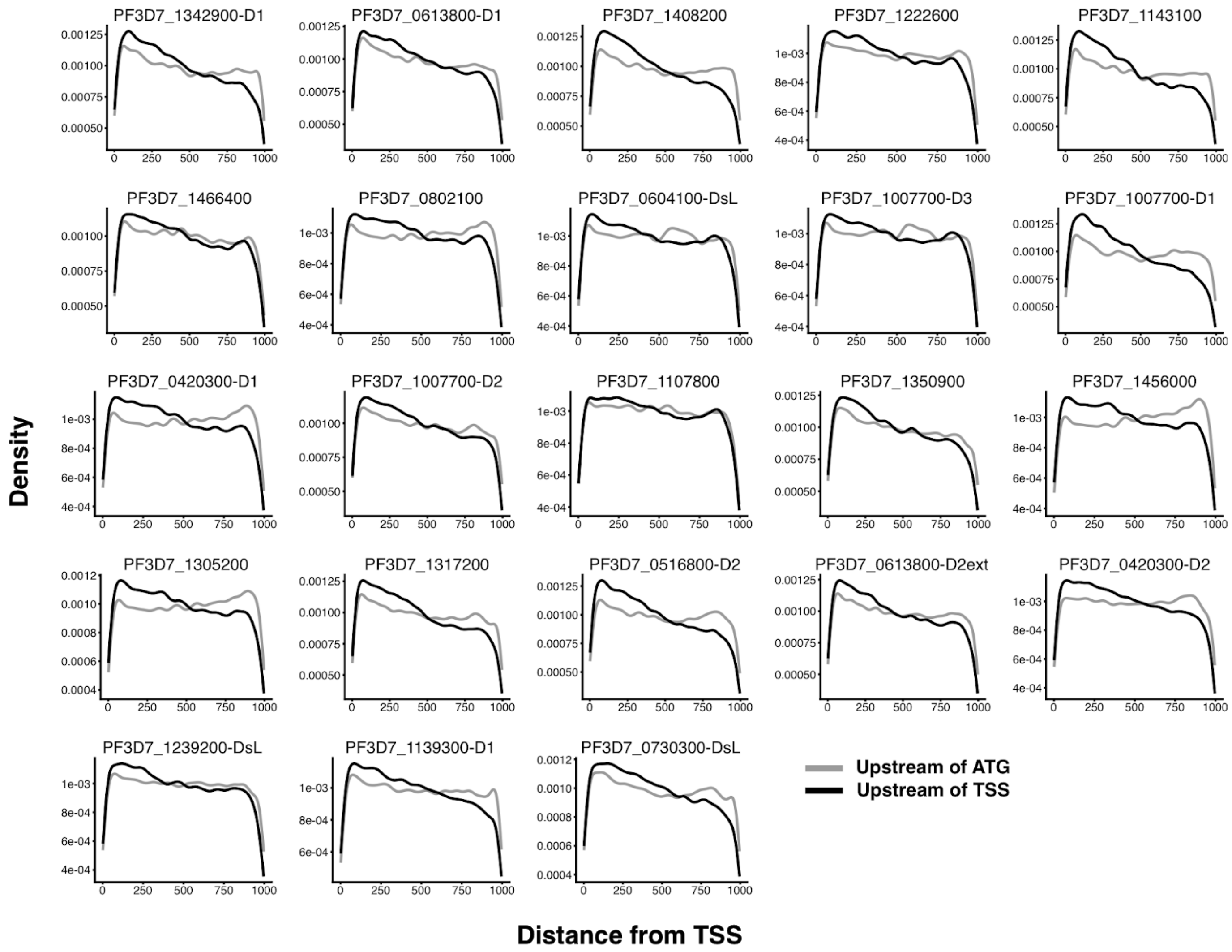


**Figure S15: ApiAP2 motif occurrences relative to predicted TSSs and annotated translation start sites in the 3D7 strain**

Motif occurrence density plots from the start (0 bp upstream) to end (1000 bp upstream) of sequences upstream of either predicted TSSs or annotated and highly curated translation start sites (ATG). We observe a consistent increase in frequency within 0–250 bp of predicted TSSs in 3D7 relative to sequences upstream of ATGs. The 3D7 strain is representative of a general trend seen in all strains.

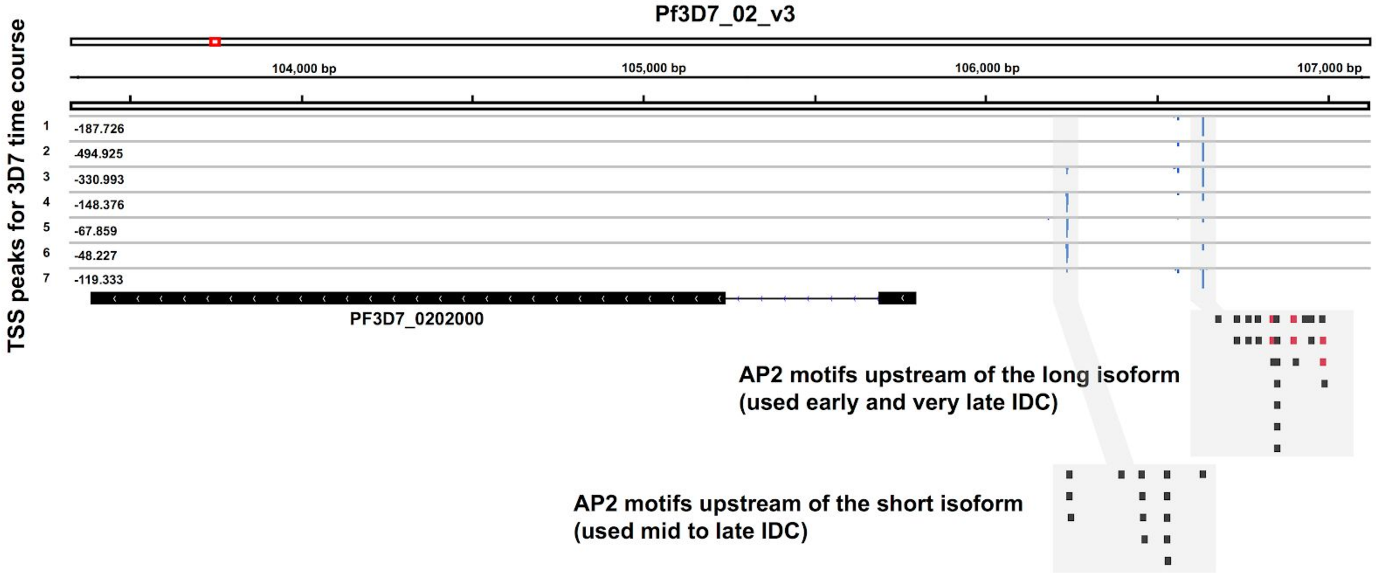


**Figure S16: ApiAP2 motifs upstream of the annotated KAHRP translation start site**

The TSS landscape of the knob-assisted histidine-rich protein (KAHRP) shows a dynamic pattern throughout the IDC and exhibits a long and short isoform of its 5' UTR. Different motif occurrences found on both strands are displayed for long and short isoforms. Motifs highlighted in red show the binding sites of the AP2 transcription factor encoded by the gene *Pf3D7_1466400* (*Pf14_0633)*: these motifs are only found upstream of the long isoform.

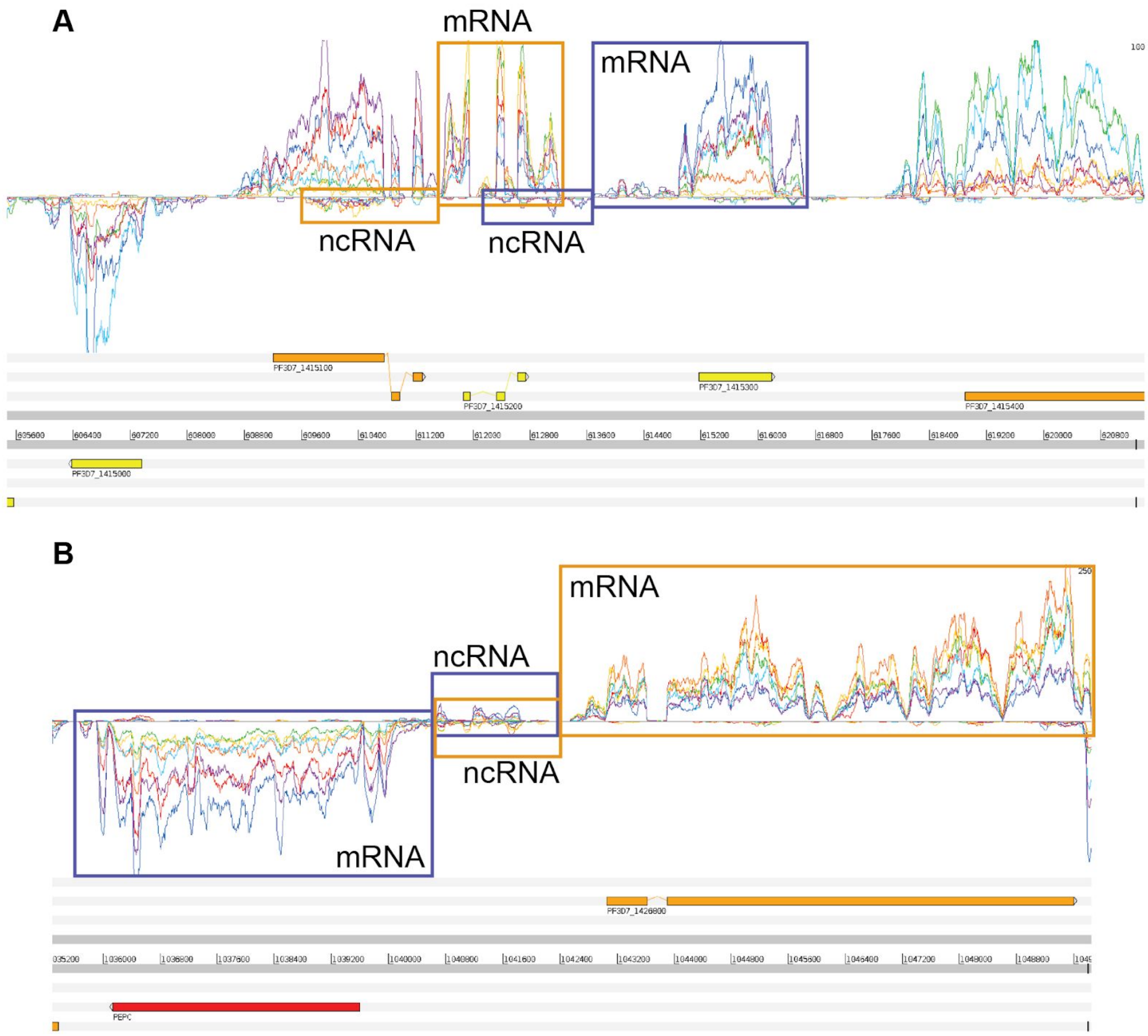


**Figure S17: TSS-associated non-coding RNAs**

Pairs of mRNA and correlated antisense non-coding RNAs can be found in a number of genomic locations, additionally to the example shown in figure 4. In panel A, one of the ncRNAs (in the purple box, bottom strand) is antisense to an upstream mRNA (orange box, top strand). In panel B, two of these ncRNAs are in antisense orientations relative to each other- information from the time enables correlated patterns of expression to define the transcript pairs.

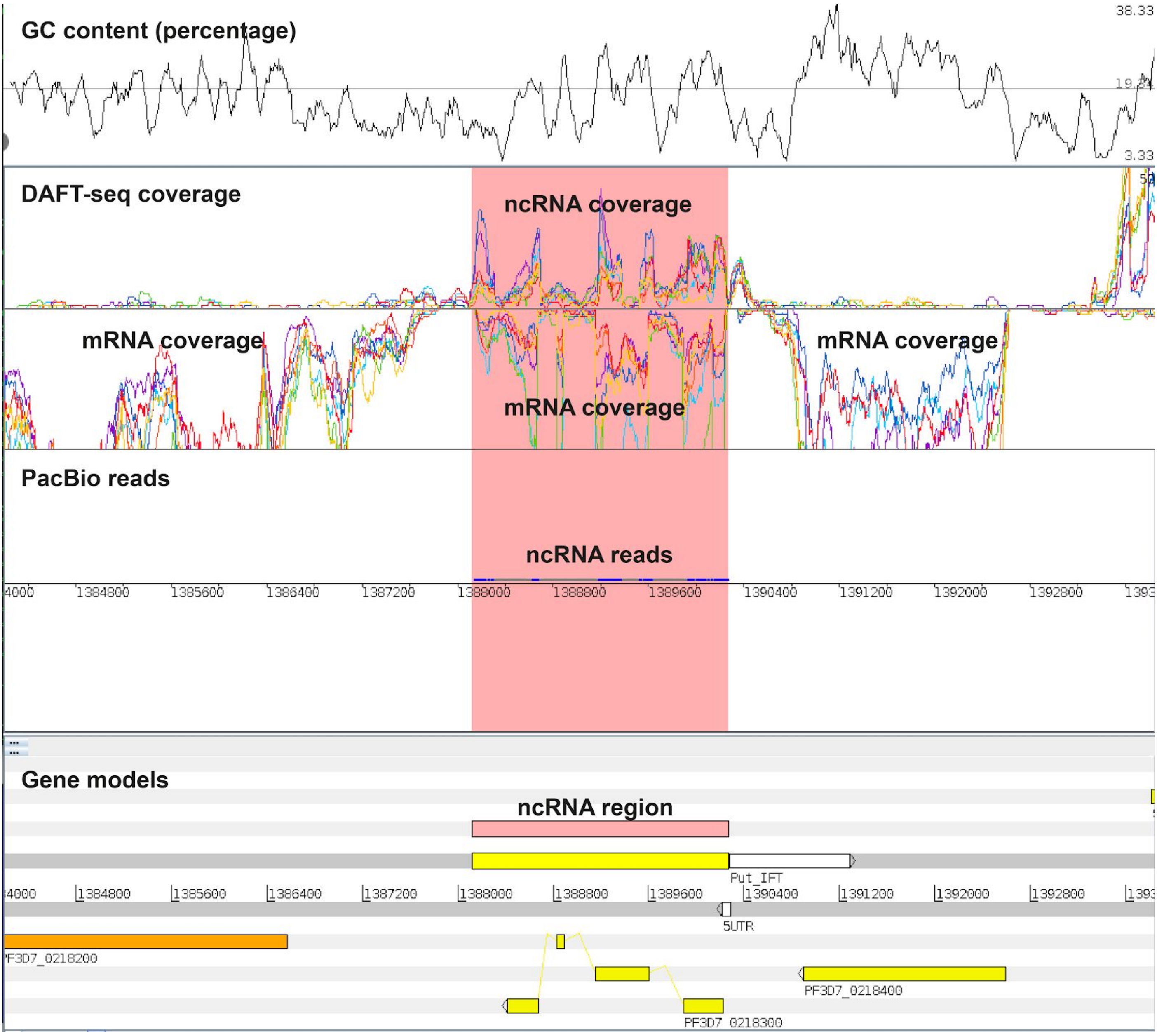


**Figure S18: A long non-coding RNA**

A non-coding RNA (present within the region highlighted in red) can be identified opposite the gene PF3D7_0218300 in both DAFT-seq and PacBio data.

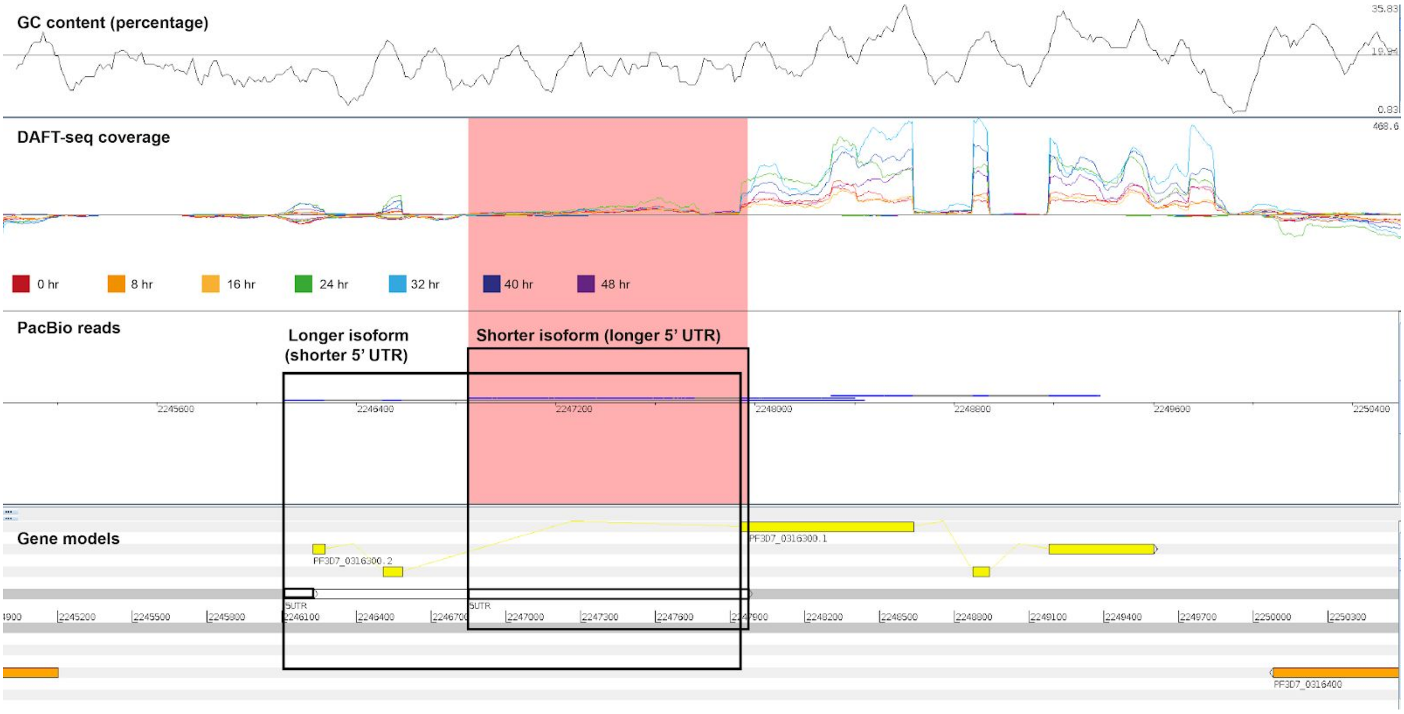


**Figure S19: Alternative splicing in PacBio reads**

Two alternative isoforms of the gene PF3D7_0316300 are captured in PacBio reads, where the reads capture different TSSs and different use of splice sites.

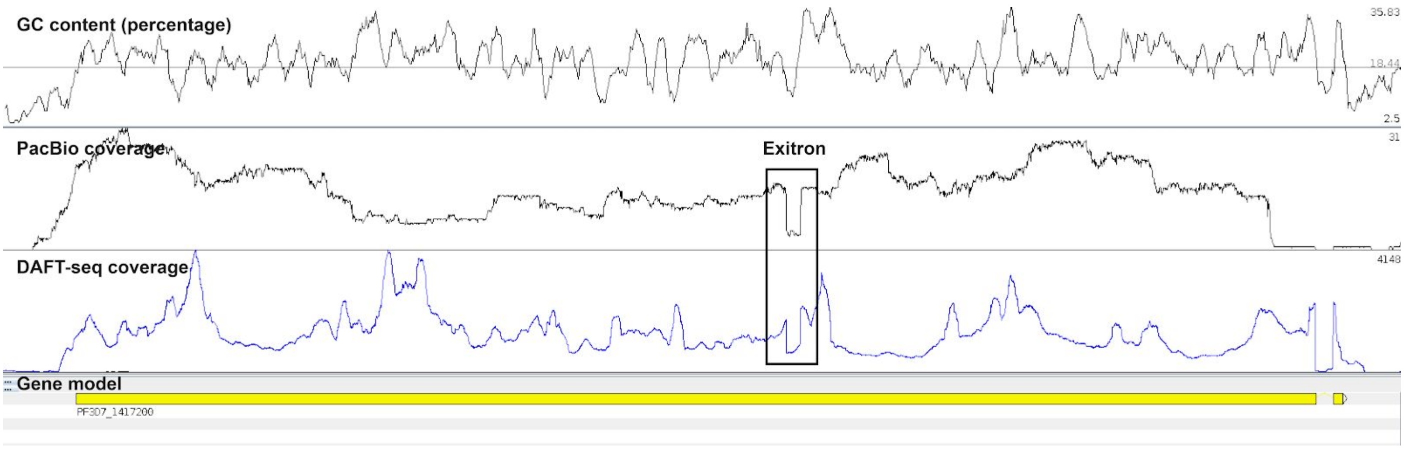


**Figure S20: Exitrons in PacBio reads**

PacBio coverage (top panel) and DAFT-seq coverage (middle panel) show the same splice site within the first protein-coding exon of the gene PF3D7_1417200, which can be identified as an exitron.

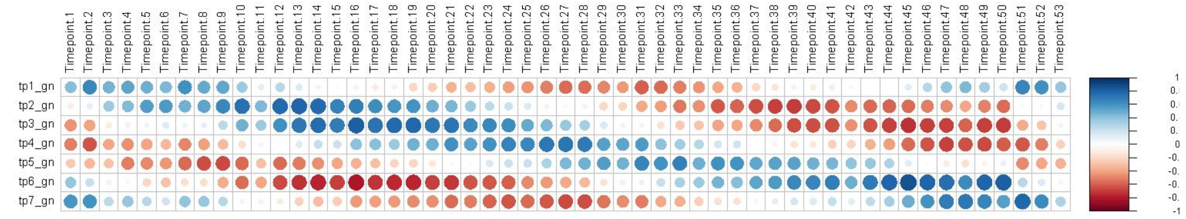


A: 3D7 DAFT-seq vs. 3D7 microarray

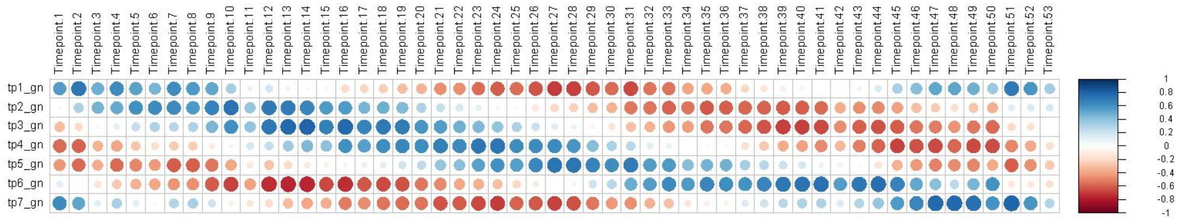


B: HB3 DAFT-seq vs. 3D7 microarray

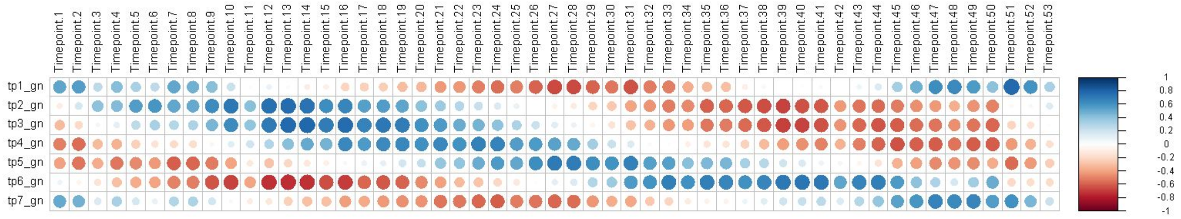


C: IT DAFT-seq vs. 3D7 microarray

**Figure S21: Correlation of DAFT-seq data to existing 3D7 microarray data by correlation plots**

Horizontal data is from a published 3D7 array time course (Llinas et al., 2006). Data on the vertical axis is from the DAFT-seq time courses, which has been normalised to better match the format of the array data (see supplemental methods). High positive correlation between two time points is shown by large dark blue circles, with lighter smaller blue circles representing weaker positive correlation. Orange circles represent negative correlation. The trend observed is that the new DAFT-seq data best matches the data in expected time point from the array data set.

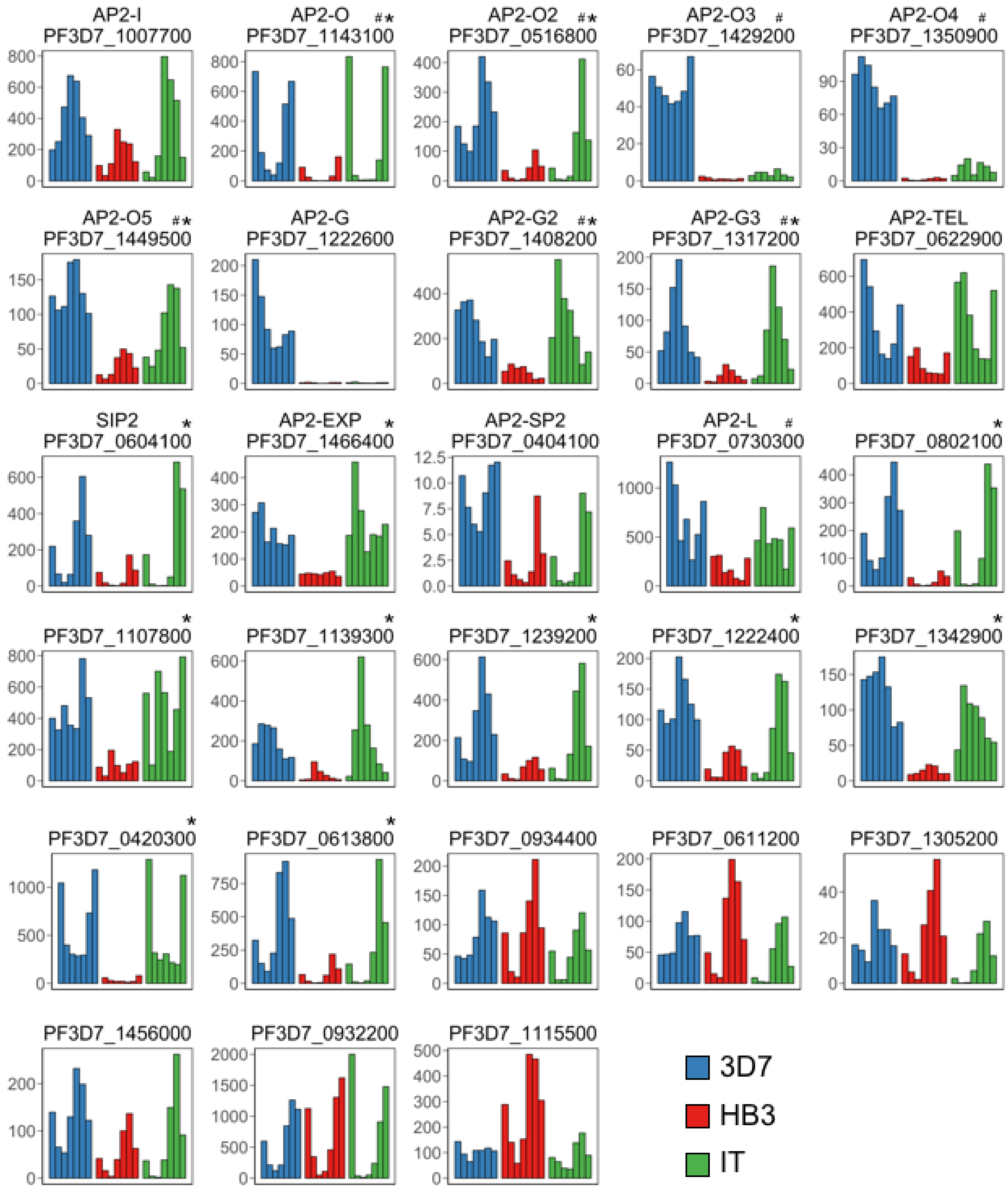


**Figure S22: ApiAP2 expression in the three *P. falciparum* strains**

Of 28 ApiAP2 transcription factors, as annotated based on the 3D7 genome on PlasmoDB, 15 are significantly overexpressed in 3D7 compared to HB3. Each plot shows the TPM calculated per time point, per strain per gene. Genes marked by an asterisk (*) are significantly overexpressed in 3D7 relative to HB3. Genes marked by a pound sign (#) represent those which have not yet been functionally validated in *P. falciparum*. All others were either not significantly different or transcription was undetected in one or both strains (TPM < 5).

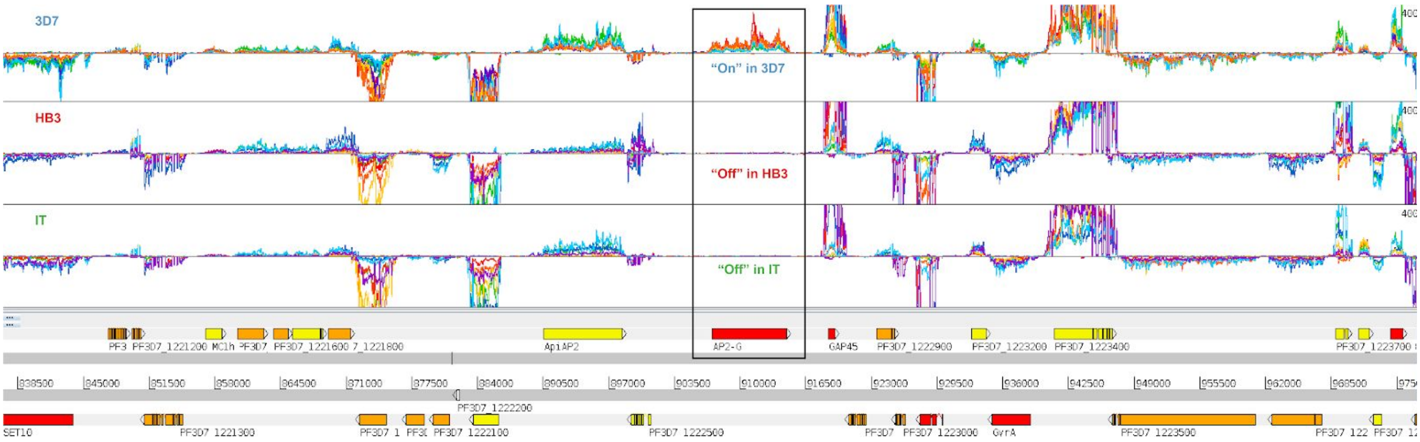


**Figure S23: AP2-G expression in the three *P. falciparum* strains**

Top panel: 3D7 DAFT-seq data; middle panel: HB3 DAFT-seq data; lower panel: IT DAFT-seq data; bottom panel: gene models (shown on both DNA strands). The seven time points are represented by the same colours in each panel (TP1=red, TP2=orange, TP3=yellow, TP4=green, TP5=light blue, TP6= dark blue, TP7=purple). Most genes show similar expression levels through the time course in each of the three strains, but the AP2-G gene is detected at higher steady state levels in the 3D7 strain (AP2-G is located within the black box).

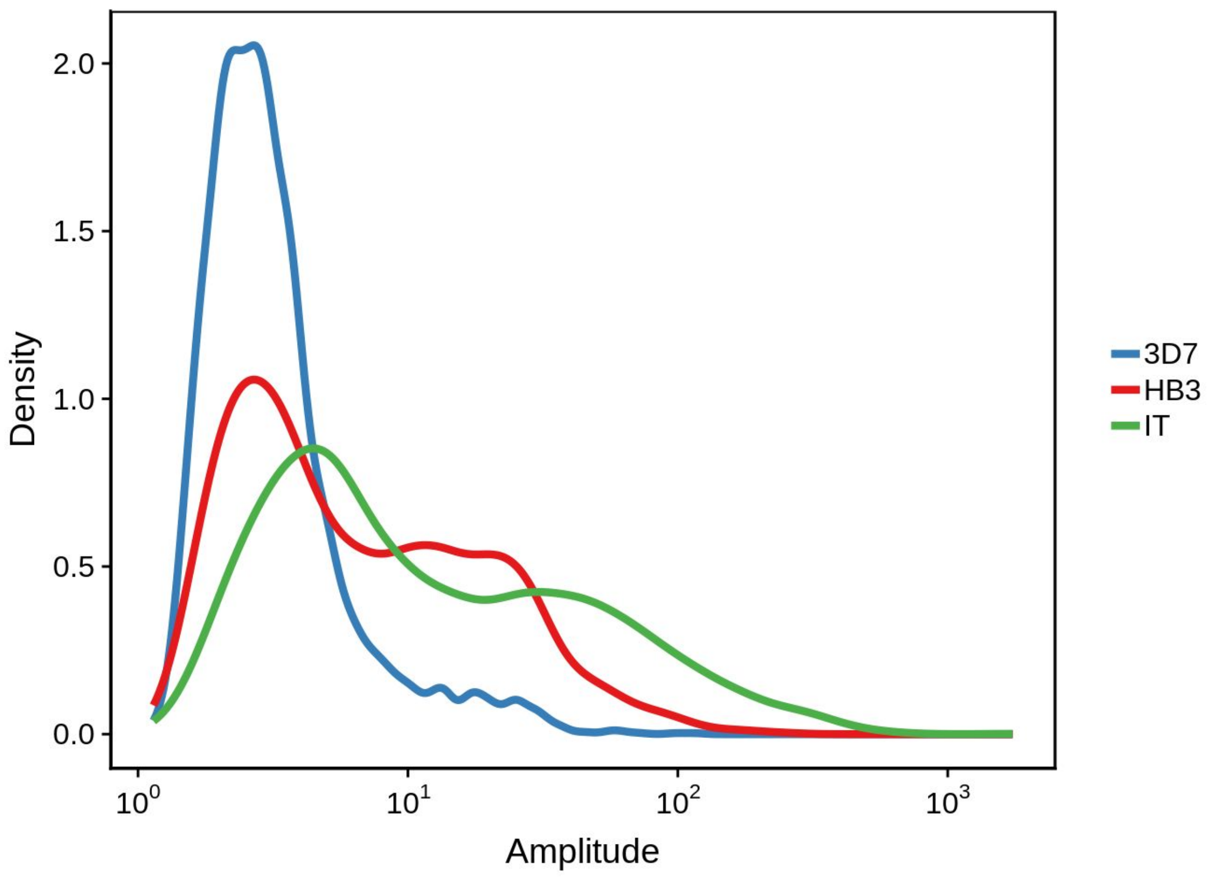


**Figure S24: Amplitude change distributions for all genes in the three *P. falciparum* strains**

Amplitudes for each gene were calculated as the difference between its maximum and minimum estimated TPM throughout the IDC.

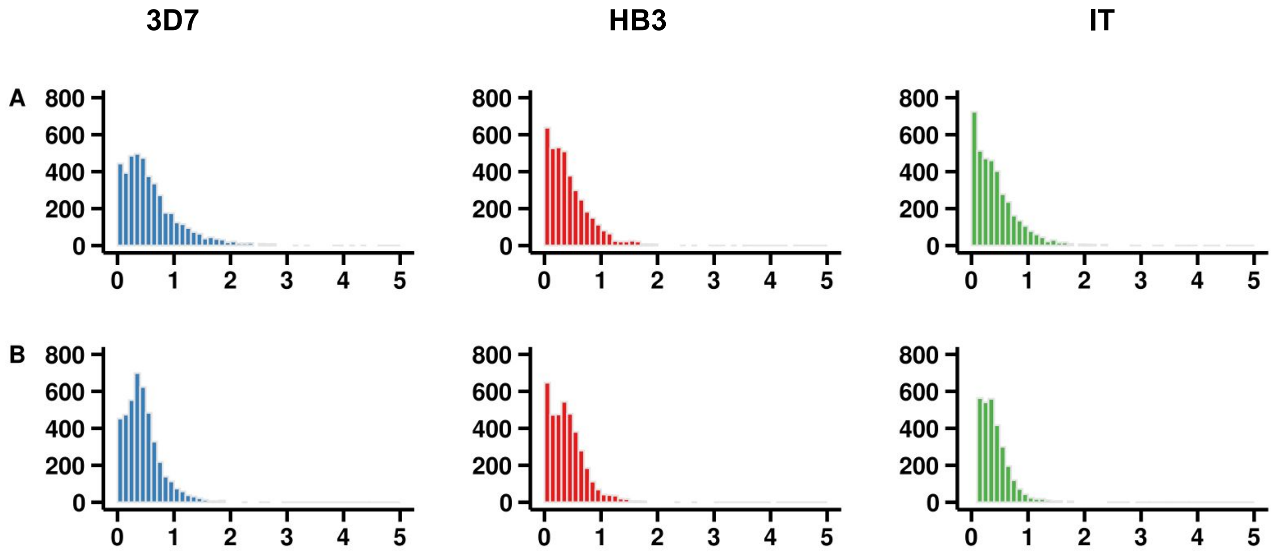


**Figure S25: Coverage-based UTRs for the three *P. falciparum* strains**

Length distributions for coverage-based UTR predictions for 3D7 (blue), HB3 (red), and IT (green) strains in kilobases (kb).

**A)** 5’ UTR length distributions. Median lengths are 476, 346, and 339, nucleotides, for 3D7, HB3, and IT, respectively.

**B)** 3’ UTR length distributions. Median lengths are 399, 356, and 287 nucleotides for 3D7, HB3, and IT, respectively.

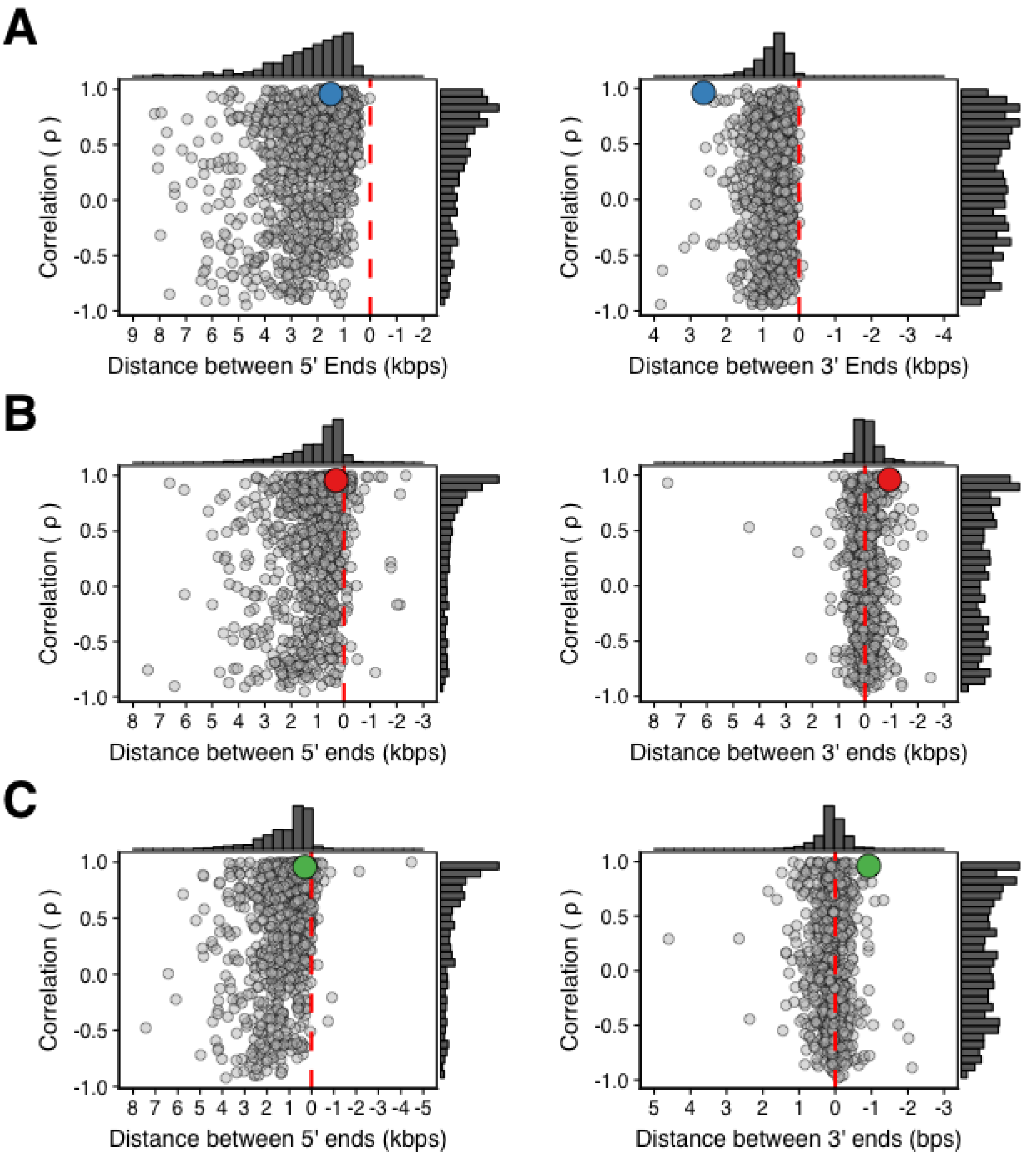


**Figure S26: Analysis of expression patterns of adjacent gene pairs for the three *P. falciparum* strains**

Correlation of gene expression (in TPM, using Pearson) for 1119 pairs of head-to-head genes and 1059 pairs of tail-to-tail genes. Large, colored data points represent the gene pair PF3D7_1011900 (heme oxygenase) and PF3D7_1012000 (putative RING zinc finger protein) in the left column and PF3D7_1115900 (DHHC9) and PF3D7_1116000 (RON4) in the right column.

**A)** Distance by correlation **without** annotated 5’ UTRs (left) or 3’ UTRs (right) plotted against the distance of intervening genomic sequence for the 3D7 strain. Without annotated 5’ UTRs, median distance between head-to-head genes is 1946 bp. Without annotated 3’ UTRs, median distance between tail-to-tail genes is 657 bp. (Note: figure 3 in main paper shows 3D7 with annotated UTRs)

**B)** Distance by correlation **with** annotated 5’ UTRs (left) or 3’ UTRs (right) plotted against the distance of intervening genomic sequence for the HB3 strain. Median distance between head-to-head and tail-to-tail genes is 713 and 23 bp, respectively.

**C)** Distance by correlation **with** annotated 5’ UTRs (left) or 3’ UTRs (right) plotted against the distance of intervening genomic sequence for the IT strain. Median distance between head-to-head and tail-to-tail genes is 762 and 70 bp, respectively.

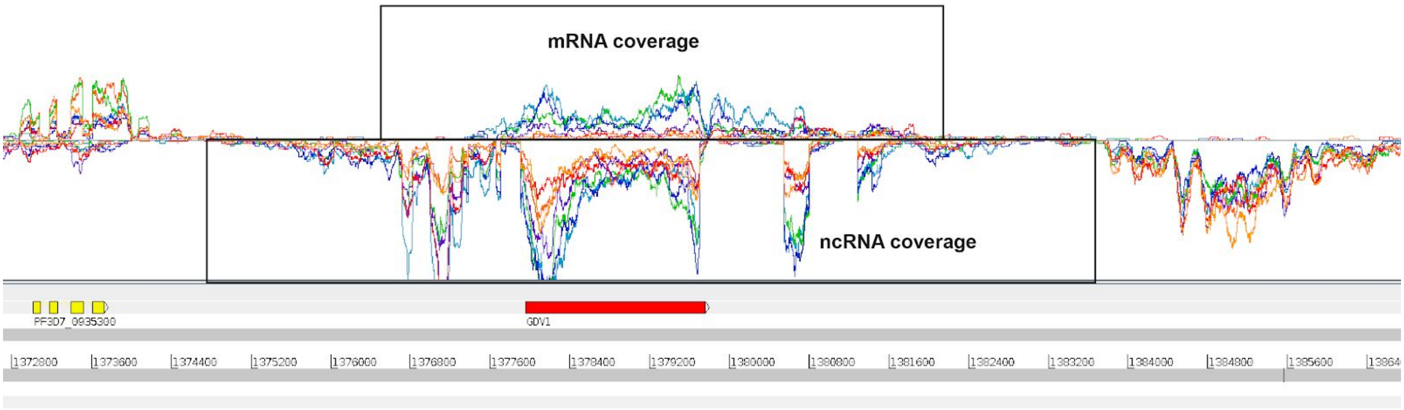


**Figure S27: A spliced non-coding RNA opposite GDV1**

A spliced non-coding RNA (marked within the region highlighted in the lower box) can be identified opposite the gene GDV1 (mRNA region marked within upper box) in the 3D7 DAFT-seq data.

### Supplementary text

#### Transcriptome realignments

We used a multidimensional scaling (MDS) based approach to analyse phase differences of peak transcript levels between strains, similar to that in (1), using the knowledge that many genes have a clear peak of expression in the IDC (2). This approach has been used in other model organisms such as yeast and humans when cyclic gene expression profiles were presumed (3, 4).

To apply MDS to our data, we first only included core, nuclear genes (i.e. non-telomeric as defined by GeneDB) that were transcribed (TPM of at least 5 within at least one sample) in all three strains. Additionally, only genes with an amplitude of at least 1.41 ($2^{0.5}$) were included in our analysis, where the amplitude was calculated as the ratio of maximum to minimum TPM values throughout the time course, in order to filter out constitutively expressed or randomly fluctuating genes.

We then applied the isoMDS function found within the MASS package (5) to each time point for each strain and generate an uncorrected time point phase diagram as seen below:

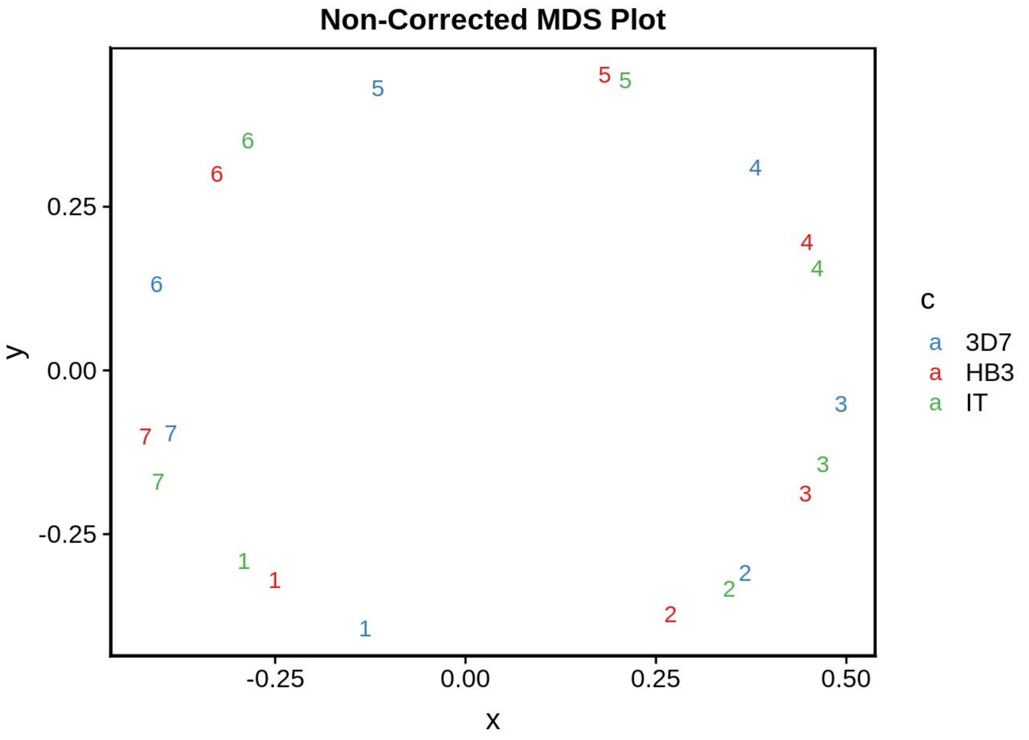


**Figure ST1: Non-Corrected MDS plot**
The colored numbers represent each of the seven time points in the three IDC time courses.

We observed that the 3D7 strain appears to be several hours “ahead” of the other strains..

We attempted to perform a local transcriptome alignment using the 3D7 53 time-point gene expression microarray from (6). The procedure was as follows:

1. TPMs were first mean normalized and standardized in order to compare to normalized microarray data. This involved defining the mean TPM value to zero. The mean and standard deviation are calculated across the time course for each gene, then subtracted from the mean for each TPM value, then divided by the standard deviation.
2. Expression values were then imputed for each hour within 48 hours for each time course by fitting a periodic spline (periodicSpline function of the splines R package, version 3.5.1, as periodic splines can interpolate a function which is periodic in nature) to the 7 observed time points, using a 48 hour cycle as an input parameter.
3. Next, a time point-by-time point correlation matrix was generated by correlating each time point within the imputed RNA-seq data set to the observed data in the 3D7 microarray data set.
4. The imputed time points within the RNA-seq data sets that were most highly correlated with a set of microarray time points(1, 8, 16, 24, 32, 40, and 48 hours) were selected. For example, if the most highly correlated time points could be 2, 7, 16, 23, 38, 47 for the imputed RNA-seq data sets. These were then interpreted as the corrected expression values for each gene at time points 1, 8, 16, 24, 32, 40, and 48.
5. Finally, another periodic spline is fit to the previous predictions in order to impute new values for the time course at hours 1, 8, 16, 24, 32, 40, and 48.
6. These expression values at time points 1, 8, 16, 24, 32, 40, and 48 were then used as input for a locally realigned MDS plot

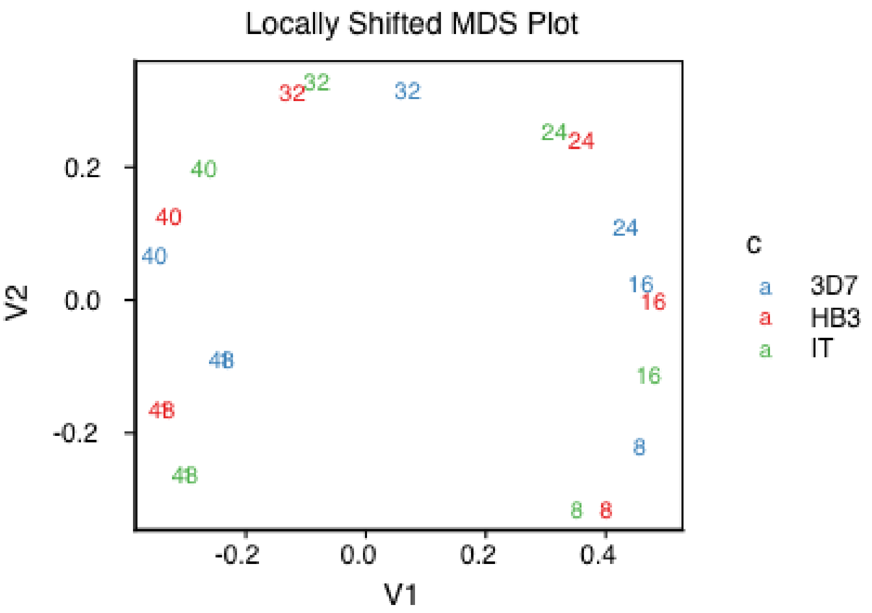


**Figure ST2: Locally shifted MDS plot**

The colored numbers represent each of the seven time points in the three IDC time courses.

We found that this method was able to successfully realign some, but not all time points. For example, time point 40 now appears to qualitatively cluster bettern, some time points did  not. Most importantly, this did not change the overall phase difference histograms when comparing strains, suggesting that correcting for this experimental artifact is likely to be computationally infeasible with current tools. As a global realignment did not result in improved phase predictions, we decided to leave uncorrected, calculated phases as is for our main analysis of comparing the phases between strains.

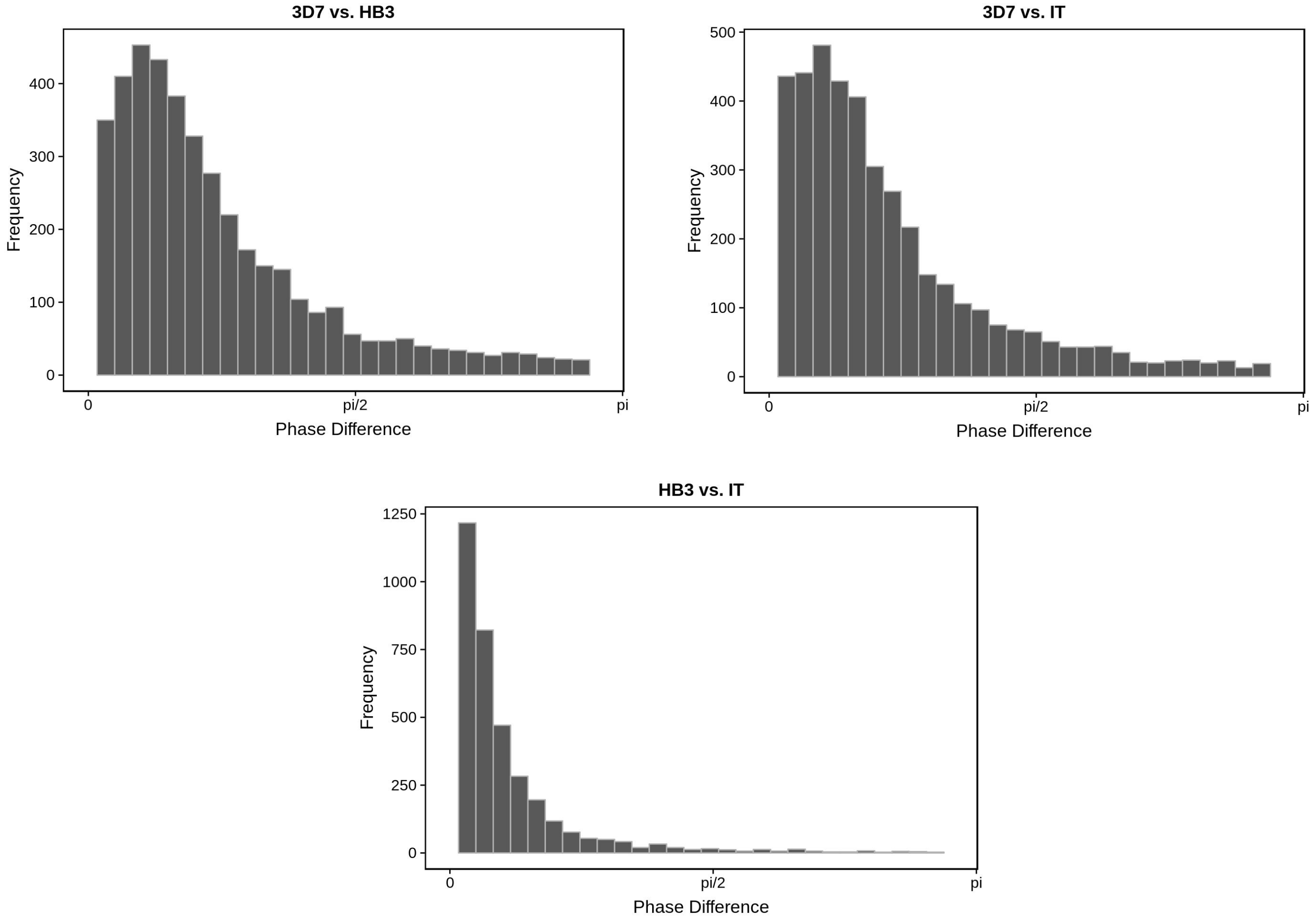


**Figure ST3: Phase differences for pairs of strains, using corrected data**

#### UTR length comparisons

We compared our UTR predictions to UTRs predicted in previous work. (7) predict 5' and 3' UTRs using RNA-seq data and a machine learning algorithm. However, they make these predictions for each time point in their time course. Similarly, (8) use a variant of CAGE sequencing (9) in order to define transcription start sites throughout the genome, for multiple time points.

For Caro et al., the longest 5' and 3' UTR predictions for each gene were used to compare to our own. Pearson correlation estimates were generated using the cor.test function in R (method="pearson").

For Adjalley et al., the longest, shortest, average, and median UTR lengths for each gene were calculated. These predictions were further filtered using a threshold cutoff of 4 as to compromise between conservative and liberal TSS predictions (see (8)) and a positive length (i.e. predicted transcription start sites cannot be downstream of annotated translation start sites). Finally, single nucleotide transcription start sites were determined by dividing the length of each "TSS block" by 2. That is, the TSS used in our comparison is the midpoint of each "TSS block."

#### Homopolymer tract analysis

We wished to confirm why coverage-based UTRs sometimes suffered from sudden drops in read coverage within blocks of otherwise continuous transcription, as this was an artifact that can artificially shortened them. We suspected that homopolymer tracts were the cause of these coverage drops. Our ability to "rescue" missing fragments of coverage-based UTRs using the 5UTR-seq (TSO-based) TSSs is consistent with this hypothesis.

Key questions include:

- Do long homopolymer tracts exist within predicted coverage UTRs?
- Are UTRs containing long homopolymer tracts on average shorter than those that don't?

To address these questions, we extended the 5' UTRs 100 bp upstream beyond their predicted coverage-based TSS (i.e. prior to 5UTR-seq assisted correction), counted the length of the longest homopolymer tract for each nucleotide, and determined whether longer homopolymer tracts tend to occur in 5' UTRs that are shorter. In order to account for the possibility that longer sequences will be more likely to contain longer homopolymer tracts by chance, the UTR lengths were normalized by the lengths of the longest homopolymer tracts found within each sequence. As is seen in Figure S9, we do find that shorter UTRs tend to have longer homopolymer tracts. Furthermore, if we focus only on UTRs that could be "repaired", the correlation gets stronger and is strongest when we look only at 100 bp upstream of "repaired" 5' UTRs. This suggests that 5' UTRs in *P. falciparum*, using short-read technology, can only be effectively predicted using a combination of RNA-seq and TSS-seq.

#### Generating Position Weight Matrices

Core motifs were identified as in (10). Full-length position weight matrices (PWMs) representing *in vitro* DNA-binding motifs for several ApiAP2 DNA-binding domains, were downloaded and manually converted to MEME format. PWMs were trimmed from the 5' and 3' ends to include only the six most informative positions (the core) of each PWM as calculated by the Shannon entropy.
